# The maize recombination landscape evolved during domestication

**DOI:** 10.1101/2024.11.04.621928

**Authors:** Ruth Epstein, JJ Wheeler, Melissa Hubisz, Qi Sun, Robert Bukowski, Jingjing Zhai, Wei-Yun Lai, Edward Buckler, Wojtek P. Pawlowski

## Abstract

Meiotic recombination is an important evolutionary process because it can increase the amount of genetic variation within populations through the breakage of unfavorable linkages and creation of novel allelic combinations. Despite the plethora of knowledge about population-level benefits of recombination and numerous theoretical studies examining how recombination rates can evolve over time, there is a lack of empirical evidence for any hypotheses that have been put forward. To alleviate this gap in knowledge, we characterized the evolution of the recombination landscape in *Zea mays* ssp. *mays* (maize) during its domestication from *Zea mays* ssp. *parviglumis* (teosinte), explored hypotheses that permitted the evolution of the maize recombination landscape and tied these alterations to changes in the genetic basis of recombination. Using experimental populations and the population genomics approach of ancestral recombination graph (ARG) inference, our data demonstrated that maize had a 12% increase in its genome-wide recombination rate during domestication. Although the maize and teosinte recombination landscapes are highly correlated, r = 0.85 at 1Mb resolution, maize has evolved to have higher recombining regions in interstitial chromosome regions, compared to teosinte which only harbors high recombining regions sub-telomerically. Our data show that the re-patterning of COs towards interstitial chromosome regions came from reduced CO interference levels within maize. Supporting the idea that CO interference is reduced within maize, we found evidence for selection acting on trans-acting recombination-modifiers that participate in the class I CO pathway or CO interference directly. Lastly, we showed that the re-patterning of COs was beneficial to maize evolution because regions that significantly increased in recombination were targeted to gene-rich regions harboring domestication related loci. Because we found regions with significant increases in recombination had a lower deleterious mutation load, compared to regions with decreases in recombination, we concluded that the domestication-related variation in these regions, in which selection acted upon during domestication, was shielded from the Hill-Robertson effect. In conclusion, the re-patterning of CO events during domestication allowed maize to adapt and evolve at a faster rate than previously understood.

## Introduction

Meiotic recombination is an important evolutionary process because it can increase and maintain the amount of genetic variation in a population, create more advantageous genotypes to increase fitness, and boost the efficiency of selection through the breakage of linkages (1, 2). These benefits at the population level are made possible by mechanisms that initiate and control recombination at the molecular level. Meiotic recombination begins with programmed double-stranded breaks (DSBs) by a complex involving Spo11 proteins and Mtop6b (3, 4). The MRN endonuclease complex, including Mre11, Rad50, and Nbs1, cleaves dsDNA and resects 3’ overhangs (5). Only a small fraction of DSBs will be resolved into crossover (CO) events (6), which generate the reciprocal exchange of genetic material between homologous chromosomes. There are two types of COs: class I and class II (7). The ZMM suite of proteins, which involves Zip proteins, Mer3, Msh4, and Msh5, protects CO intermediates allowing the Mlh1-Mlh3 complex to resolve intermediates into class I COs (8). Class I COs are subject to CO interference—which occurs when a CO begins to form too closely to an adjacent CO, and the CO intermediate dissolves (7). The class II COs are not subject to interference, make up only ∼15% of COs per meiosis, and is controlled by the Mus81 endonuclease (9).

Although the mechanism of recombination is generally conserved (10), recombination rates can still evolve over time. Theoretical studies have shown that in order for an increase in recombination to be favorable, negative linkage disequilibrium (LD), defined as advantageous alleles appearing in the same haplotype less often than expected, must be generated throughout the population (11, 12). More recombination is needed to separate these unfavorable linkages thus bringing about conditions in which an increase in recombination would be extremely beneficial. Previously, an analysis using cytological data from 46 domesticated vascular plant species, found that the wild progenitors of domestication species had fewer COs per meiosis, than their domesticated pair, demonstrating that plant domestication events often result in higher recombination rates (13). Intuitively, since most domestication events involve directional selection resulting in a genetic bottleneck, this reduces the effective population size (N_e_) allowing genetic drift to become strong generating negative LD throughout the population (14, 15) thus allowing favorable conditions for recombination rates to increase.

There is a great body of theoretical studies that have put forward numerous hypotheses explaining how recombination rates can evolve over time (16–20). However, most of these hypotheses lack any empirical support and furthermore, studies that have experimentally characterized recombination landscape evolution during crop domestication events, are often not tied back to theoretical frameworks (21, 22). To address this gap in knowledge, we simultaneously investigated the evolution of the recombination landscape in *Zea mays* ssp. *mays* from two perspectives: a population genomics perspective, exploring hypotheses that enabled recombination landscape evolution, and a molecular perspective, examining how the loci controlling recombination were altered during domestication. We utilized the well-documented history of maize which was domesticated from *Zea mays* ssp. *parviglumis* (teosinte) by early agriculturists in Mexico about 9,000 years ago prompting maize to undergo a radical phenotypic change (23, 24). Using the population genomic method of ancestral recombination graph (ARG) inference, we inferred saturated recombination maps which permitted us to study the variation between the maize and teosinte recombination landscapes. Our data show maize displays more recombination in regions closer to the centromere through the re-positioning of COs. We concluded that indirect selection acting on trans-acting genes involved in CO interference, such as *Mlh1, Mlh3*, and *Rec8*, led to more COs in gene-dense regions, thereby protecting these regions from the Hill-Robertson effect.

## Results

### The maize genome-wide recombination rate increased during domestication

To test the hypothesis that the genome-wide recombination rate in maize increased during domestication, CO events were extracted from experimental populations of maize and teosinte, by identifying parental haplotype breakpoints in the progeny. We utilized the maize nested association mapping (NAM) recombinant inbred line population, which was generated by mating 25 diverse inbreds crossed to B73, with ∼4500 progeny (25) and a population of intercrossed Palmar Chico *parviglumis* teosintes, originally collected in Palmar Chico, Mexico, resulting in ∼3000 F2 individuals (26, 27). We filtered out individuals with CO intervals greater than 100kb, because we cannot accurately map where the CO occurred within wide CO intervals, and individuals with a large number of CO events due to alignment errors being detected as CO events. Common algorithms to detect CO events rely on switching between parental haplotypes, and alignment errors resulting in spurious haplotype changes can result in false CO detection. We also normalized recombination rates by population size to accurately compare the maize and teosinte populations.

We found maize has a higher genome-wide recombination rate (Table 1), 0.707 cM/Mb, compared to teosinte, 0.621 cM/Mb, a 12% increase. Teosinte has a longer genome length due to more knobs (28), however, all the teosinte sequences have been previously mapped to the B73v4 reference genome (29), so the physical map length is identical between maize and teosinte. This is relevant because there are more false positive COs in teosinte due to reads being mis-aligned to the B73 assembly. Although this initial analysis confirmed that maize recombination rates did increase during domestication, the number of recombination events per population was too small to make finer comparisons between the two populations.

**Table 1.**
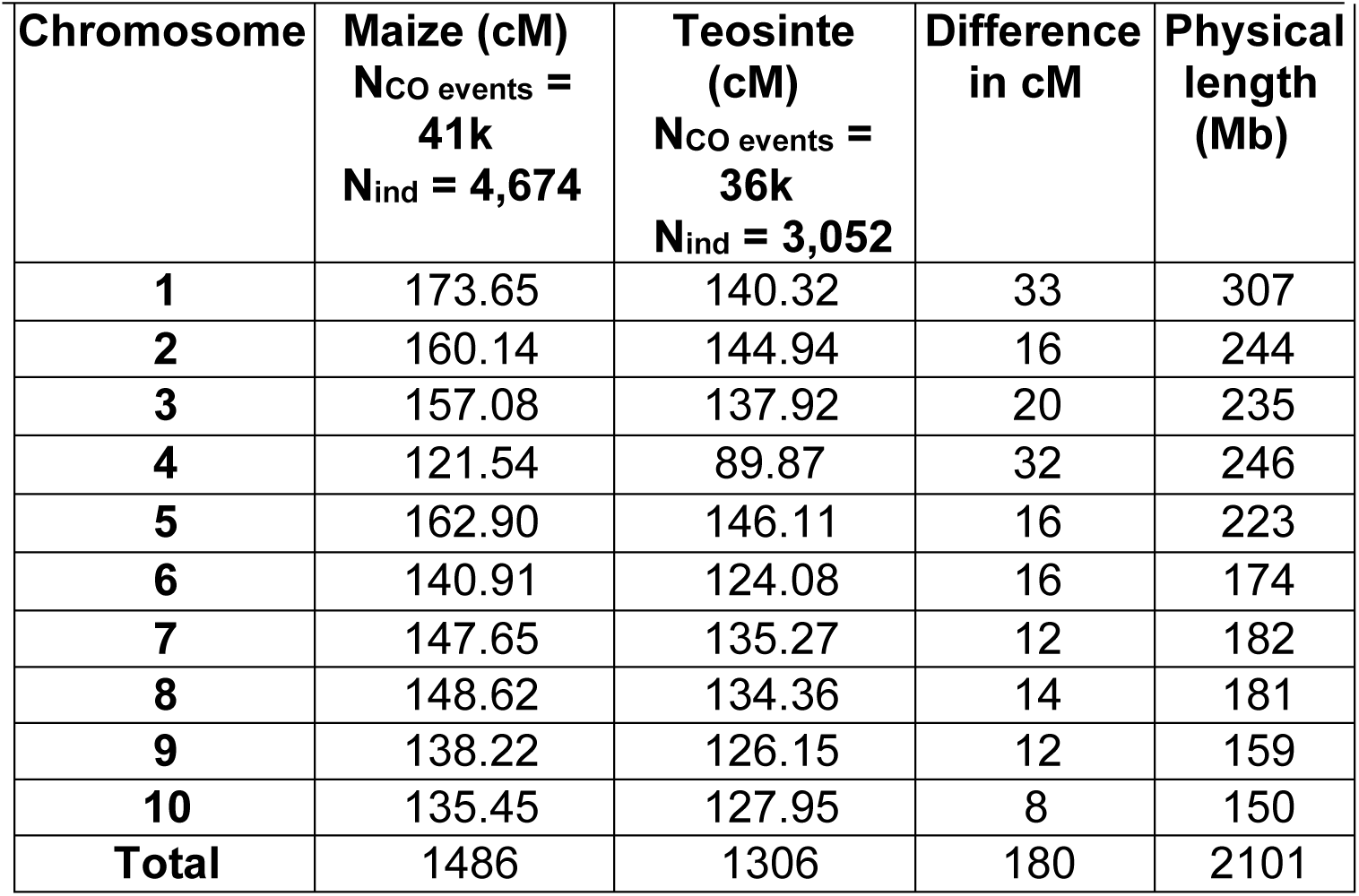
Genetic map lengths of maize and teosinte by chromosome and their differences. COs were included if they were less than 100kb. Chromosomes 1 through 5 are considered “long chromosomes” while chromosomes 6 through 10 are considered to be short.

| **Chromosome** | **Maize (cM)**<br>**N<sub>CO</sub> events =**<br>**41k**<br>**N<sub>ind</sub> = 4,674** | **Teosinte (cM)**<br>**N<sub>CO</sub> events =**<br>**36k**<br>**N<sub>ind</sub> = 3,052** | **Difference in cM** | **Physical length (Mb)** |
| --- | --- | --- | --- | --- |
| **1** | 173.65 | 140.32 | 33 | 307 |
| **2** | 160.14 | 144.94 | 16 | 244 |
| **3** | 157.08 | 137.92 | 20 | 235 |
| **4** | 121.54 | 89.87 | 32 | 246 |
| **5** | 162.90 | 146.11 | 16 | 223 |
| **6** | 140.91 | 124.08 | 16 | 174 |
| **7** | 147.65 | 135.27 | 12 | 182 |
| **8** | 148.62 | 134.36 | 14 | 181 |
| **9** | 138.22 | 126.15 | 12 | 159 |
| **10** | 135.45 | 127.95 | 8 | 150 |
| **Total** | 1486 | 1306 | 180 | 2101 |

### Maize and teosinte have similar recombination landscapes but differ in placement of high recombining regions

In order to dissect the differences between the maize and teosinte recombination landscapes at a finer scale, we needed a saturated recombination map for each population. Traditional methods of identifying recombination events to obtain a saturated map would require a large experimental population, which is expensive and resource intensive. Instead, we inferred historical recombination events utilizing ancestral recombination graph (ARG) reconstruction. To do this, we used a maize population consisting of 60 temperate and tropical inbred lines (Supp. Table 4) and a teosinte population consisting of 50 teosinte individuals sampled from 5 different locations in Mexico (Supp. Table 5) (30, 31). The ARG delineates the complete evolutionary history of a population and describes all coalescence events, which trace lineages into common ancestors, and recombination events, which split lineages (32, 33). After convergence of the ARGweaver algorithm, we extracted recombination rates by counting the number of recombination events by branch lengths from both ARGs at a single time point (see Materials & Methods for details).

We found that the patterns of maize and teosinte recombination rates across chromosomes are highly correlated with each other at the 1Mb resolution (Pearson correlation, r = 0.851) and at the 100kb resolution (r = 0.744). Both of these populations show the typical U-shaped distribution of COs, seen in most large crop genomes, in which most CO events occur towards chromosome ends with a large suppression region around the centromere and the pericentromeric region (Figure 1, Supp. Figure 1).

**Figure 1.**
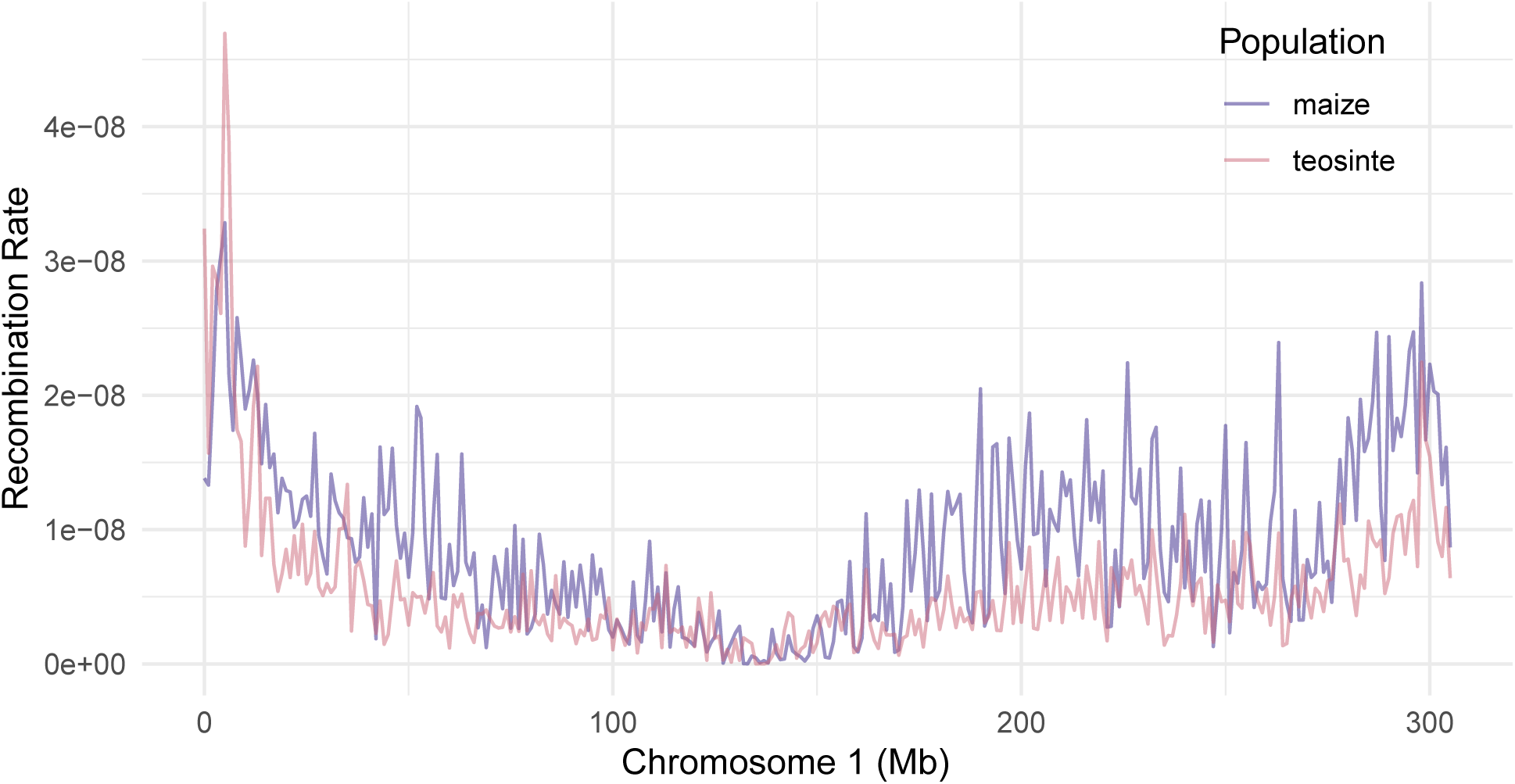
ARG-inferred maize and teosinte recombination landscape of chromosome 1.

Although the ARG-inferred map has many more recombination events than any previous genetic map created in maize, we validated this new dataset by comparing the ARG-inferred recombination map to previously identified CO hotspots from the NAM population (553 CO hotspot intervals, mean length = 15kb). We found the average ARG-inferred recombination rate at NAM CO hotspots were 2 standard deviations above the mean ARG-inferred recombination rate demonstrating that the ARG was able to accurately identify high recombining regions. In further support of our findings, 85% of NAM CO hotspots were contained in the top 15% of genome-wide recombination rate intervals.

We used a 1Mb resolution in our analyses because random fluctuations in recombination rates create noise when only 50 to 60 individuals are used per ARG. ARG construction involves a trade-off: at a 1Mb resolution, the recombination signal greatly exceeds the noise, and it is more computationally efficient. Using more than 60 individuals would drastically increase the computation time and would make inference of the true ARG more difficult.

High recombining regions in maize are interesting to note since they typically harbor genes, more genetic variation, and a lower mutation deleterious load compared to lower recombining regions (25, 34). Thus, we wanted to assess if teosinte has a different placement of high recombining regions than maize. We divided the maize and teosinte genomes into 1Mb intervals and defined the top 15% of intervals with the highest CO rate as high recombining regions (Table 2). We found that maize and teosinte do not share about 30% of their high recombining regions. We conducted this experiment for the top 5% and top 10% of high recombining regions and results were similar amongst all classes. In both populations, high recombining regions are located sub-telomerically; however, maize has more high recombining regions occurring interstitially on the longest chromosomes (Figure 2, Supp. Figure 2). We defined interstitial chromosome regions to be the midpoint between the centromere and closest telomere or regions that are closer to the centromere than the telomere.

**Figure 2.**
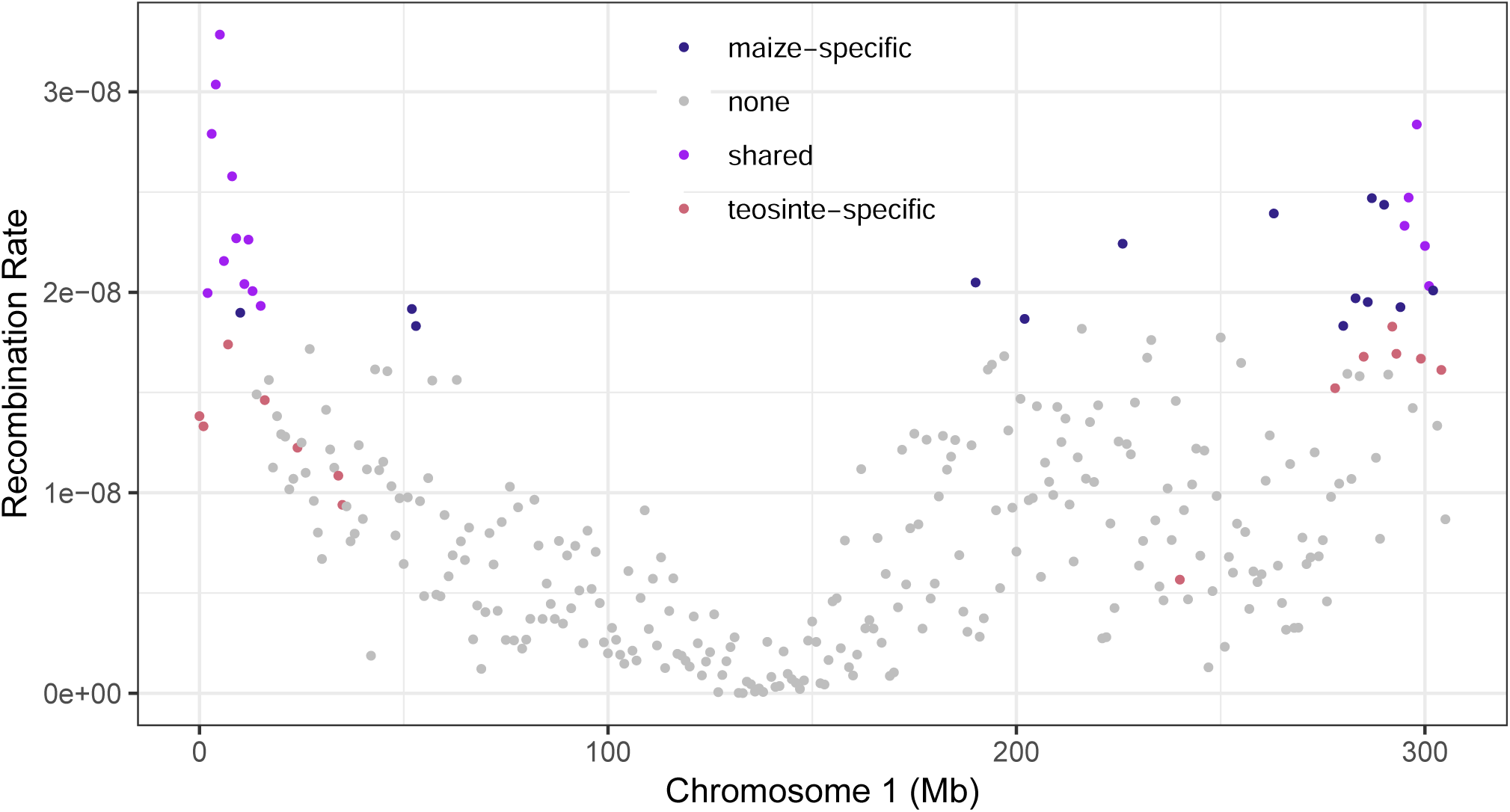
Chromosome 1 with high recombining regions highlighted. Blue = maize high recombining regions, Pink = teosinte high recombining regions, and Purple = shared high recombining regions.

**Table 2.**
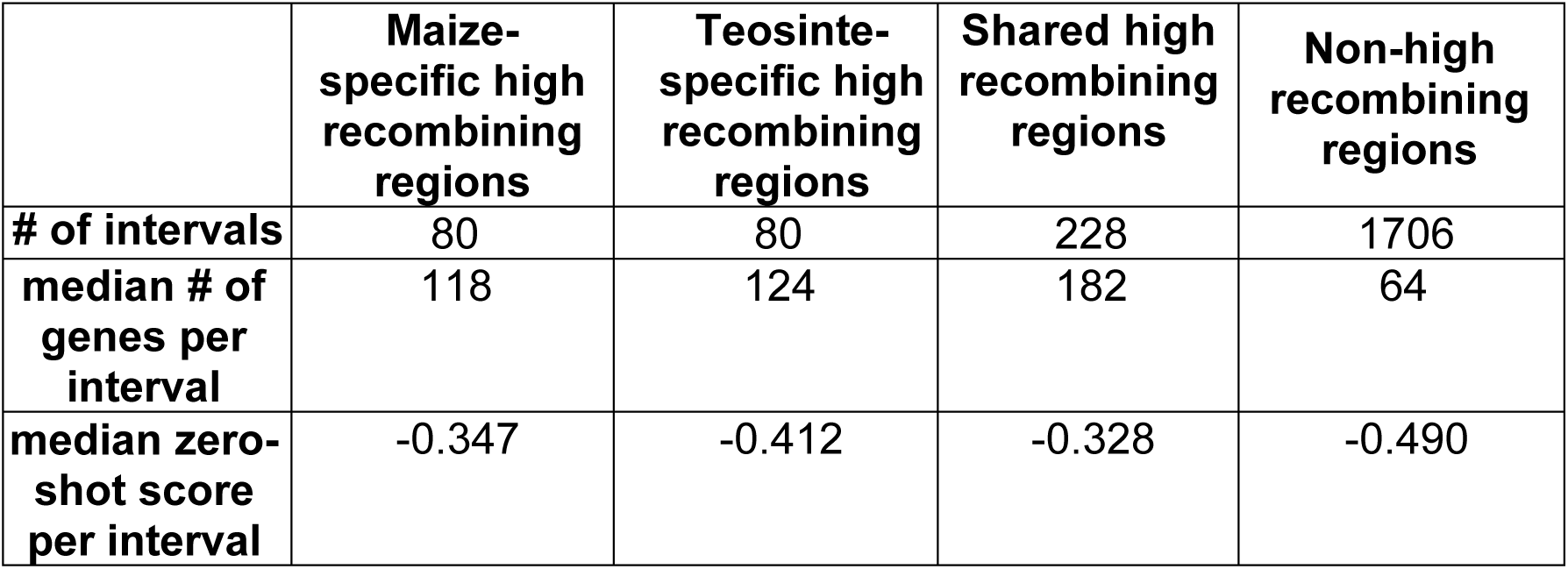
Description of the top 15% highest recombining 1Mb intervals specific to maize, specific to teosinte, and those shared between the 2 populations. Gene density counts based on B73v4 annotation. Zero-shot scores are based on the P(ref/alt), so a more negative zero-shot score signifies a more deleterious site.

|  | **Maize-specific high recombining regions** | **Teosinte-specific high recombining regions** | **Shared high recombining regions** | **Non-high recombining regions** |
| --- | --- | --- | --- | --- |
| **# of intervals** | 80 | 80 | 228 | 1706 |
| **median # of genes per interval** | 118 | 124 | 182 | 64 |
| **median zero-shot score per interval** | -0.347 | -0.412 | -0.328 | -0.490 |

We then investigated if high recombining regions that are not shared between maize and teosinte were responsible for the genome-wide increase of the recombination rate in maize. We termed regions with significantly different recombination rates between maize and teosinte, differentially recombining regions or DRRs. We considered a region a DRR if the interval was 2 standard deviations above or below the mean difference between the maize and teosinte recombination rates. High recombining regions specific to maize are not the same as maize DRRs because maize DRRs can still include regions with low recombination rates, as long as they are greater than the teosinte rate.

There were more than twice as many maize DRRs as teosinte DRRs (Table 3). Similar to the high recombining region analysis, maize DRRs were concentrated interstitially on chromosomes, much closer to the pericentromeric region, while teosinte DRRs were found mostly in sub-telomeric regions (Figure 3, Supp. Figure 3).

**Figure 3.**
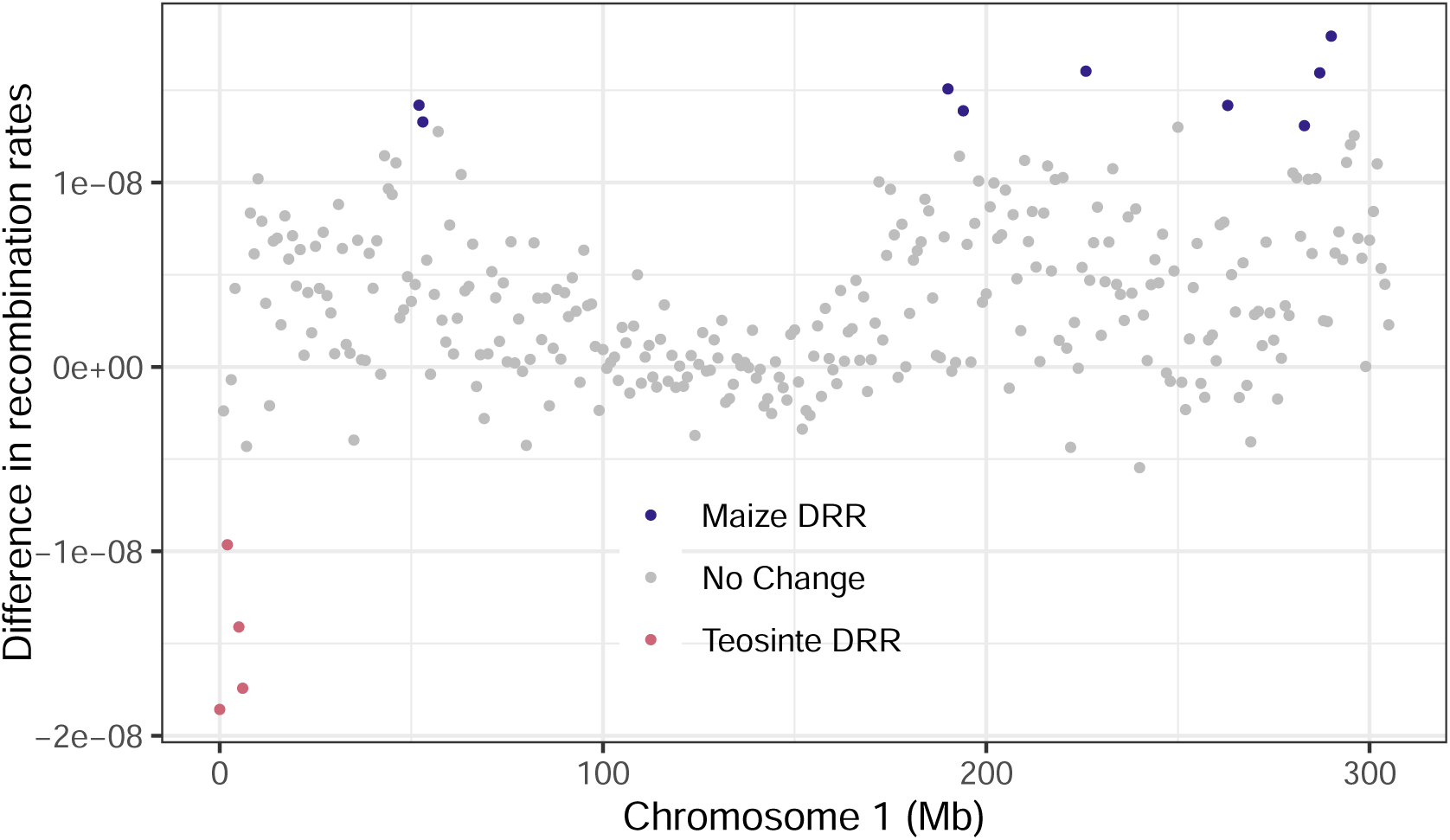
Maize and Teosinte DRRs on chromosome 1. Blue = maize DRR, Pink = teosinte DRR, grey = less than 2 standard deviations away from mean difference.

**Table 3.** Description of DRRs in maize and teosinte genome wide. DRRs are 2 σ away from the mean difference between maize and teosinte recombination rate per chromosome.

|  | **# of 1Mb intervals** | **# in sub-telomeric regions** | **# in interstitial regions** |
| --- | --- | --- | --- |
| **Maize DRRs** | 74 | 51 | 23 |
| **Teosinte DRRs** | 31 | 27 | 4 |

### The increase of recombination rates in maize was towards more gene-dense regions

For recombination to be beneficial, it should occur in or near genes to create advantageous allelic combinations. Thus, we wanted to assess if the observed increase in recombination during domestication was advantageous through investigating if maize COs are more biased towards gene dense regions compared to teosinte. Using the B73v4 genome annotation, containing about 40,000 genes, and the ARG-inferred recombination landscapes, we found the maize recombination landscape, at a 1Mb resolution, and gene density were more correlated (Pearson’s correlation, r = 0.652) than the teosinte recombination landscape (r= 0.617). At a 100kb resolution, the maize recombination landscape was still more correlated with gene density patterns (r = 0.576) than teosinte (r = 0.524). The difference in correlation demonstrates the regions in maize that have significantly increased in recombination, are skewed toward more gene dense regions to ensure recombination occurs within genes creating more meaningful allelic combinations. We also assessed if regions that had substantial increases in recombination in maize harbored large-effect loci that were selected during domestication, but we did not find any such associations.

Another advantage of recombination is its ability to purge deleterious mutations from populations. Previous studies have shown maize recombination hotspots have a lower deleterious load compared to cold spots, or lower recombining regions (25). We then hypothesized that regions that became highly recombining in maize during domestication would have reduced deleterious load compared to regions that experienced decreases in recombination. The deleterious mutation dataset was constructed using a zero-shot score strategy which hypothesizes that mutations maintained in conserved coding sequences are likely to be deleterious (36). The score of how deleterious a mutation is, was based on the log-likelihood differences between the reference and alternative allele, where a more negative value is a more conserved deleterious site. The dataset was trained on a diverse panel of *Zea*, and zero-shot scores were predicted at ∼25M sites using the 26 NAM parents and SNPs found in repeat or transposon-rich regions were removed. We considered the top 15% of maize and teosinte recombination rates at a 1Mb resolution to be high recombining regions, and selected intervals that were unique to each population.

The median genome-wide zero-shot score was –0.490. The median zero-shot score at maize-specific high recombining regions was –0.347, while the median zero-shot score at teosinte-specific high recombining regions was –0.412 (Table 2). A t-test between these two groups showed there was a significant difference in the means (t = 3.8169, df = 148.66, p-value = 0.0001978) demonstrating that deleterious load is more depleted at regions that became high recombining in maize during domestication. Nevertheless, teosinte-specific high recombining regions still had a lower deleterious load than genome-wide because these regions have maintained a recombination rate higher than the genome-wide average in maize.

### Maize has reduced CO interference compared to teosinte

Finding that the longest maize chromosomes had an increase in recombination rates interstitially with a small decrease sub-telomerically, led us to investigate if the distribution of CO events across a chromosome was altered during domestication. The genetic map lengths of maize and teosinte chromosomes, based on the experimental population mentioned above, indicate that the longest chromosomes in maize experienced the most significant increases in genetic map lengths (Table 1). Similarly, the ARG-inferred dataset showed the longest maize chromosomes contain more DRRs enriched interstitially than sub-telomerically (Figure 3, Supp. Figures 20-29), while the shortest chromosomes show fewer interstitial DRRs. Based on these observations, we hypothesized that CO interference on the longest chromosomes was reduced in maize.

We then explored if the degree of CO interference was altered during domestication. Since the ARG and the experimental mapping populations cannot track which COs occurred in the same meiosis, we used how far each CO midpoint is from the nearest telomere as a proxy to measure the degree of CO interference. Although interference can move across the centromere (37), the pericentromeric region of maize is so devoid of COs, it is very unlikely that two COs would emerge in this region and interfere with each other. Furthermore, a previous study involving a diverse panel of maize inbred lines found a negative correlation between the level of CO interference and the distance to the nearest telomere (38). Maize chromosomes do not have a centromere in the exact middle of the chromosome, which creates a long arm and short arm of each chromosome, with chromosome 6 being the most extreme. We considered chromosomes 1 through 5 to be long and chromosomes 6 to 10 to be short. Using ∼41,000 COs in maize and ∼36,000 COs in teosinte from the initial experimental populations, we found that genome-wide, maize COs on long chromosome arms were about 14.8Mb from the telomere, while teosinte COs were only 9.8Mb from the telomere (Table 4). Conversely, the average CO distance to the telomere on short chromosome arms were quite similar between maize and teosinte, 6.9Mb and 6.3Mb, respectively. Supporting the hypothesis that long chromosome arms and the longest chromosomes had the largest alterations in CO interference levels, the shorter the chromosome arm or the shorter the entire chromosome, the more similar the maize and teosinte CO distances to the telomere are.

**Table 4.** Distance of CO midpoint to telomere on 10 long and short chromosome arms based on the B73v4 genome length.

| **Chromosome # (L=long arm, S=short arm)** | **Maize median CO distance to nearest telomere (Mb)** | **Teosinte median CO distance to nearest telomere (Mb)** |
| --- | --- | --- |
| **1S** | 16.26 | 12.09 |
| **1L** | 47.4 | 23.94 |
| **2S** | 10.68 | 9.52 |
| **2L** | 24.47 | 14.7 |
| **3S** | 4.942 | 3.93 |
| **3L** | 25.62 | 19.26 |
| **4S** | 6.90 | 6.10 |
| **4L** | 39.678 | 7.01 |
| **5S** | 6.946 | 6.68 |
| **5L** | 12.05 | 9.94 |
| **6S** | 6.18 | 4.44 |
| **6L** | 14.47 | 11.53 |
| **7S** | 5.26 | 5.02 |
| **7L** | 15.29 | 9.64 |
| **8S** | 10.6 | 10.4 |
| **8L** | 9.25 | 6.678 |
| **9S** | 12.45 | 13.12 |
| **9L** | 9.26 | 7.677 |
| **10S** | 4.27 | 4.31 |
| **10L** | 8.00 | 6.682 |
| **Median short arm dist.** | **6.924** | **6.394** |
| **Median long arm dist.** | **14.88** | **9.796** |
| **Median dist.** | **12.8** | **9.3** |

### Changes in DNA methylation are not responsible for the differences in the CO landscape between teosinte and maize

Several studies investigating the predictive power of the recombination landscape have shown epigenetic factors such as low cytosine methylation, histone modifications such as H3K4me3, and high nucleosome occupancy to be the main predictors (39, 40). Arguably, all of the abovementioned features create more accessible chromatin and therefore these features can be summarized by a single predictor, usually DNA methylation, to explain how likely a region is to undergo recombination (39, 41). Thus, we explored if DNA methylation alone was responsible for the creation of maize DRRs. Even though the maize and teosinte methylomes were not significantly different from each other, 5278 differentially methylated regions (DMRs) were previously detected (42), suggesting de-methylation during domestication could have given rise to the changes in the maize recombination landscape. DMRs were detected if there was a 40% difference in methylation levels, for each methylation context, between maize and teosinte (42).

As DNA methylation and CO location have a negative relationship, we hypothesized that if DNA methylation was involved in altering recombination, maize DRRs would be enriched in maize hypo-DMRs. However, we found this not to be the case, maize hypo-DMRs were not enriched in maize DRRs, and teosinte hypo-DMRs were not enriched in teosinte DRRs.

We then hypothesized that DNA methylation changes that altered the recombination landscape were smaller than 40%. To test this hypothesis, we trained a model to predict recombination rates using maize and teosinte-specific DNA methylation patterns. We compared the levels of DNA methylation to recombination rates in 1Mb interval using high-quality cytosine methylation data, ^m^CG and ^m^CHG, extracted from maize and teosinte individuals (42). We did not use ^m^CHH data because the majority of CHH sites were unmethylated across maize and teosinte populations (42). Although the DNA methylation data came from leaf tissue extracted from the third leaf stage, previous studies have shown that at CO sites the differences in methylation patterns between leaf tissue and meiotic tissue were negligeable (43). The genome was binned into the same 1Mb segments as mentioned above, each region was annotated with the maize and teosinte ARG-inferred recombination rate and the mean DNA methylation level in both ^m^CG and ^m^CHG contexts. We then constructed an ordinary least squares linear regression to predict recombination rates.

The maize-specific model showed ^m^CG explains 70% of variation in maize recombination rates (mean R^2^ = 0.70 ± 0.04, mean RMSE = 0.269 ± 0.0217, Supp. Table 1). Since ^m^CG is highly correlated with ^m^CHG (r > 0.95), it was not included in the model and using ^m^CHG instead of ^m^CG slightly reduced R^2^ by 0.1. Next, we constructed the teosinte-specific model and found that teosinte-specific DNA methylation could explain 65% of the variation in the teosinte recombination landscape (mean R^2^= 0.65 ± 0.04, mean RMSE = 0.256 ± 0.0183, Supp. Table 2). ^m^CHG was a slightly better predictor than ^m^CG because ^m^CG reduced R^2^ to 0.63.

We hypothesized that if teosinte DNA methylation patterns can explain a similar amount of variation in the maize recombination landscape as it does in the teosinte landscape, this suggests DNA methylation changes did not alter CO patterns in maize. We found that the maize model explains 64% of the variation in the teosinte recombination landscape compared to the teosinte model explaining 65% (R^2^ = 0.64, RMSE = 0.0682, Supp. Figure 4) and the teosinte model explains 67% of the variation in the maize recombination landscape while the maize model explains 70% (R^2^ = 0.68, RMSE =0.2760 Supp. Figure 5).

Despite differences between the maize and teosinte recombination landscapes, the teosinte model could explain more variation in the maize recombination landscape than in teosinte, demonstrating that alterations in DNA methylation levels during domestication were not related to the changes observed in the maize recombination landscape. Instead, we found that regions that display increased recombination in maize already had low DNA methylation levels in teosinte. We found the average maize-specific ^m^CG and ^m^CHG methylation levels at maize DRRs was 75% and 61% respectively, while the average teosinte-specific ^m^CG and ^m^CHG methylation levels at the same maize DRRs was 74% and 62% respectively.

### Evidence of selection on trans-acting recombination-modifiers

Since we determined that DNA methylation, did not play a role in altering the maize recombination landscape during domestication, we hypothesized that trans-acting factors, genes involved in the progression of recombination, might be involved. Several studies have shown there is heritable recombination rate variation within maize, meaning there is variation in the loci controlling recombination (38, 44), termed recombination-modifiers. Previous hypotheses suggest recombination can evolve in one of two ways: indirect selection from the hitchhiking of modifiers with other selected loci or direct selection acting on modifiers.

To assess if hitchhiking caused the genetic basis of recombination to evolve in maize, we investigated if modifiers were in domestication selective sweeps. Selective sweeps are regions experiencing very strong selection that show severely reduced genetic variation around the selected locus and loci linked to it. We used a previously constructed domestication selective sweep dataset based on a cross-population composite likelihood ratio (XP-CLR) approach (45) between teosinte and maize individuals. We then assembled a list of about 120 recombination-related genes in maize using the maize genome annotation and homology to recombination genes identified in the Arabidopsis reference genome (Supp. Table 3). We then inspected if domestication selective sweeps contain any of the identified recombination-modifiers.

We found multiple modifiers detected within domestication sweeps including *Spo11-2*, *Exo1A*, *Prd2*, *Rad50*, *Pms1*, *Mlh3*, *Figl1*, *Mtop6b1*, and *Afd1* (Figure 4). *Spo11-2*, *Prd2*, and *Mtop6b1* are involved in the formation of programmed DSBs (3, 46, 47), *Exo1A* facilitates the resolution of CO intermediates into COs (48), *Rad50* is a part of the MRN complex cleaving dsDNA and processing DSB overhangs (5), *Mlh3* is a Mut-L homolog and is directly involved in class I CO resolution (49), *Pms1* is also a Mut-L homolog but is involved in mismatch repair (50), *Figl1* encodes a pro-CO factor in the class I CO pathway in maize (51), and *Afd1,* a homolog of *Rec8,* is a part of the chromosome axis and the meiotic sister-chromatid cohesion complex (52–54). Supporting these findings, a previous study showed *Spo11-2* to have a slightly higher level of genetic diversity in teosinte compared to maize, indicative of it being subject to selection during domestication (55). Although it is possible that these genes could be the target of the selective sweep, it is unlikely that modifiers were under strong enough selection to cause a sweep, making it more likely that they were swept due to physical linkage with other selected loci, since the targets of most domestication selective sweeps are related to reproductive or environmental response that were under strong selection during domestication (26, 56).

**Figure 4.**
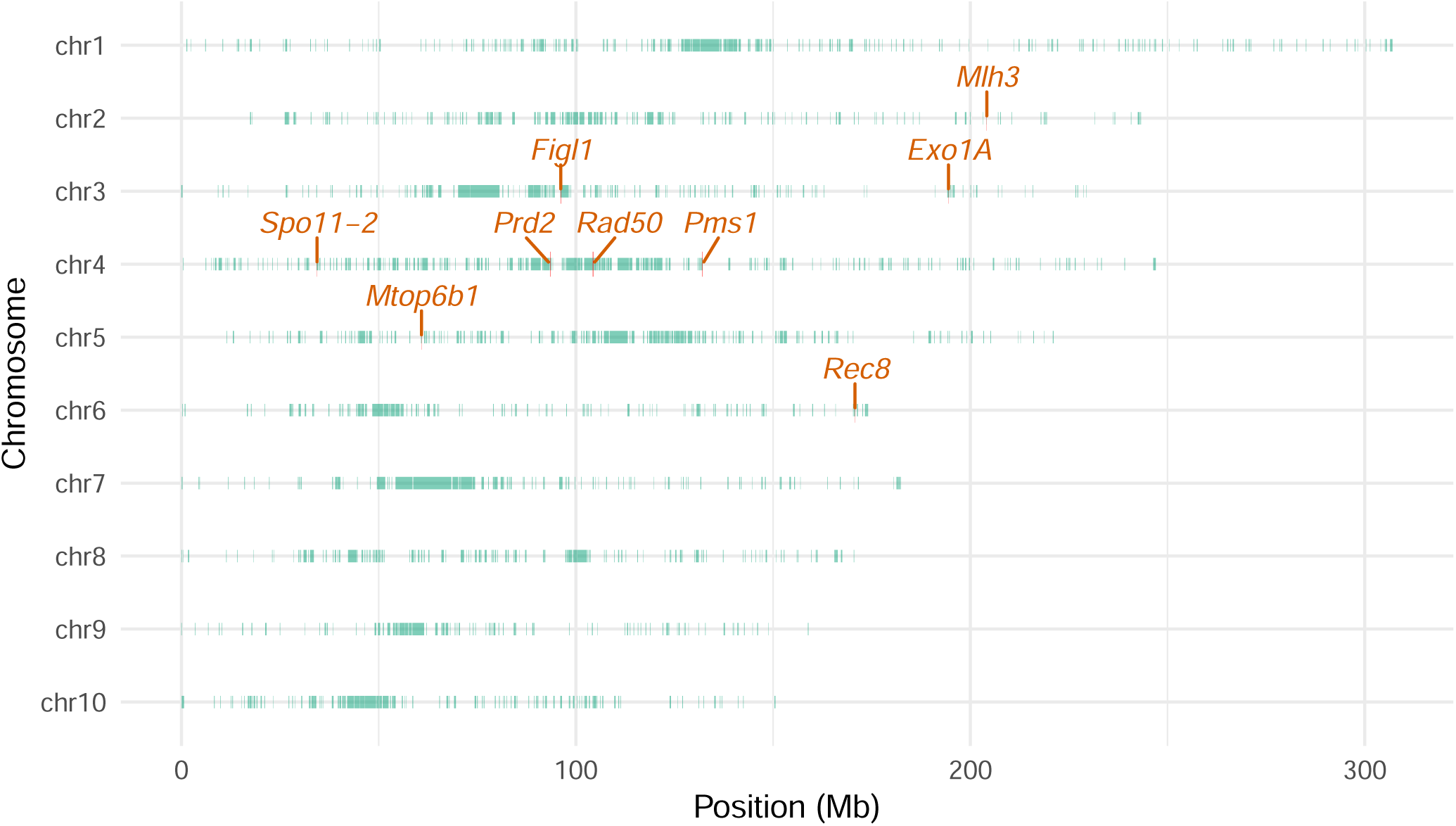
Previously identified domestication selective sweeps annotated with recombination modifiers that overlap sweeps.

Since modifiers in sweeps were subject to mostly indirect selection, we decided to quantify any selection acting on modifiers during maize domestication. As the ARG contained the complete evolutionary history of our population of interest, it was used to investigate selection signatures throughout the genome. RTH values – the relative half-time to the time to the most recent common ancestor (TMRCA), which describes the ratio between the time when half of lineages coalesce and the TMRCA, were used as a selection signature. It has been shown that particularly low values of RTH signal a clustering of coalescence events suggesting a recent partial selective sweep (32). RTH values are particularly interesting because they rely on the topology of the ARG rather than overall rate of coalescence meaning RTH values depends on time in the ARG rather than the probability of coalescence. However, RTH values can only detect partial selective sweeps, not completed sweeps.

Thus, RTH values were used to detect deviations that might have been missed in a large genome-wide selective scan and were supplemented with an F_ST_ analysis, which explain the level of genetic differentiation at a locus regardless of if a selective sweep is complete or not. After LD pruning to reduce spurious F_ST_ estimates between tightly linked SNPs, F_ST_ was calculated, as the proportion of the genetic variation within maize and teosinte relative to the total genetic variance between maize and teosinte, using 776k SNPs genome-wide shared between the 60 maize inbred lines and the 50 teosinte individuals.

We found that the mean RTH value genome-wide was 0.533. Using our previously constructed list of modifiers, *Exo1A*, *Rec8*, *Mlh3*, and *Mlh1* all displayed RTH values less than 0.2 (Table 5). Interestingly, *Exo1A*, *Rec8*, and *Mlh3* were found to be in domestication selective sweeps but were on the edges of the sweep regions, explaining why they have low RTH values. *Mlh1*, which was the only gene with a low RTH value not in a sweep, forms a complex with *Mlh3* (49) and was previously shown to have signatures of positive selection during maize domestication (55).

**Table 5.** RTH and FST values of recombination-modifiers in sweeps or with low RTH values.

| **Gene name** | **RTH value** | **F<sub>ST</sub> value** |
| --- | --- | --- |
| *Spo11-2* | 0.45 | 0.09 |
| *Exo1A* | 0.14 | 0.6 |
| *Rad50* | 1 | 0.1 |
| *Pms1* | 0.25 | 0.24 |
| *Mlh3* | 0.054 | 0.15 |
| *Figl1* | 0.96 | 0.5 |
| *Rec8* | 0.122 | 0.26 |
| *Mlh1* | 0.127 | 0.23 |

The F_ST_ analysis yielded a genome-wide median F_ST_ of 0.0648, which is similar to previous estimates of differentiation between maize and teosinte (45). Out of the original list of modifiers, *Exo1A*, *Pms1*, *Mlh3*, *Figl1*, *Rec8*, and *Mlh1* had the highest F_ST_ values, suggesting their nucleotide composition were significantly altered during domestication (Table 3). Although we identified some of these genes above because they were in domestication sweeps, these genes could have been homogeneous and non-polymorphic before domestication, so being swept with other selected loci would not result in any allele differences. Interestingly, all these genes have roles in class I CO formation, including CO interference. Compared to other genes under selection during domestication (27), these F_ST_ values were relatively low, due to these modifiers being under purifying selection reducing the ability for significant genetic differences to emerge (10).

## Discussion

### Maize recombination landscape evolved via reduced CO interference via trans-acting modifiers

We found that the maize recombination landscape evolved during domestication, resulting in more COs interstitially, with a small trade-off of fewer COs sub-telomerically on the longest chromosomes. Our data show a reduction in the level of CO interference within maize was associated with a re-patterning of CO events across chromosomes, specifically on long chromosome arms. Mechanistically, adding an extra CO per meiosis to a long chromosome arm is easy and can be achieved at the cost of only a slight reduction in interference. In contrast, shorter chromosomes and short chromosome arms did not experience as significant an increase in recombination during domestication, likely due to interference being more tightly regulated on shorter chromosomes (47).

Our data show that while DNA methylation is a key predictor of CO location in plants, it was not responsible for altering maize recombination rates. Given the importance of DNA methylation in controlling gene expression and transposable element activity (57, 58), there are limits to how much DNA methylation levels can be modified. For instance, maize mutants with more than a 20% reduction in DNA methylation are often embryo lethal (59). Other cis-acting factors, like structural variation, can influence recombination landscapes (16). The main difference between the maize and teosinte genomes is the presence of knobs which have been shown to both increase and suppress local recombination rates (28, 60, 61). However, we cannot investigate how the presence of teosinte-specific knobs impact recombination since all the teosinte data has been aligned to the B73 genome assembly. Although knobs can impact recombination, they have no impact on nearby genes and LD structure, which suggests their effect is probably minimal (60) and that the genome-wide changes we observe in recombination rates during domestication are due to other phenomena. Structural variation aside, the general decrease in any genetic variation during the domestication of maize could have led to more recombination, since there is a greater level of sequence similarity between homologous chromosomes (62). However, the level of polymorphisms often effects recombination locally, not globally which we cannot detect given our analysis is at 1Mb resolution (39, 62, 63), and furthermore, investigating the effect of polymorphisms on recombination would be confounding given ARG creation relies upon SNPs to infer the genealogical history of a given population.

Instead, we propose that trans-acting factors were responsible for altering the maize recombination landscape during domestication. Modifying genes involved in the class I CO pathway and CO interference would only alter recombination, could occur rather quickly, and could be less costly than altering DNA methylation. Since COs were merely repositioned and the maize methylome was not altered significantly during domestication, regions where recombination increased were already in open chromatin, and therefore no changes in DNA methylation were necessary.

The recombination-modifiers we identified to be under selection, *Exo1A, Mlh3, Mlh1, Figl1*, and *Rec8,* play important roles in modulating CO positioning on chromosomes through participation in either interference-dependent class I CO formation or CO interference directly. Interestingly, targeting CO interference seems to be a common mechanism to alter CO positioning and genome-wide recombination rates. For example, in an experimental evolution study of Drosophila, there was evidence for indirect selection reducing the level of CO interference, sometimes resulting in increased recombination or merely just the repositioning of COs (64). *Rec8*, which was detected in our selection scan, was identified as a candidate QTL influencing the novel patterning of COs, resulting in less recombination in interstitial chromosome regions, in domesticated barley (65) and has been implicated in modulating CO interference to ensure proper chromosome pairing in tetraploids (66). Although *Hei10* dosage was proposed to be responsible for the level of CO interference in Arabidopsis (8), we did not identify *Hei10* as a selection target in any of our selection scans.

We identified several modifiers to be under selection and most likely, these modifiers evolved via indirect, rather than direct selection, given most selective sweeps contain genes under strong selection previously implicated in the radical phenotypic change maize underwent during domestication (27). In further support of indirect selection being the cause of recombination evolution in maize, simulation studies have shown modifiers can be favored indirectly through strong selection acting on other loci nearby, causing recombination rates to increase (16). This idea has empirical support: studies in *Drosophila*, barley, and tomato have shown strong selection on other traits, such as those relating to domestication and local adaptation, can unintentionally cause recombination landscapes to be altered over large time scales (21, 22, 67).

### The increase in recombination during domestication was beneficial to maize evolution

Our data show that the re-patterning of CO events during maize domestication was thoroughly beneficial. Previous studies have demonstrated that polygenic domestication-related traits are controlled by large quantities of QTLs with diminishing small effects (68) indicating they are ubiquitous across the genome. The fact that the re-patterning of COs directed maize DRRs towards more gene-rich regions suggests that these regions most likely contained more domestication-related variation, in which selection acted upon. Supporting this claim, the same study compared the genetic architecture of 18 polygenic domestication-related traits and found regions with the highest recombination rates harbored a disproportionately large amount of heritable additive genetic variation before and after domestication (σ_A_) (68). This analysis showed that the maize landrace population contained more σ_A_ (85% of all σ_A_) in high recombining regions compared to the teosinte population (82.5% of all σ_A_). This observation suggests that genomic regions involved in polygenic adaptation during the domestication process became high-recombining, thereby reducing the impact of the Hill-Robertson effect (2). Evidence for this protective effect is seen in the significantly lower deleterious load found in regions of the maize genome that experienced increased recombination during domestication compared to regions where recombination rates decreased. The re-patterning and increase of COs enabled selection to act more efficiently on domestication-related variation, allowing maize to adapt and evolve at a faster rate during domestication than previously understood.

### Evolution of recombination rates has a tight upper bound

Our data demonstrate that maize had a 12% genome-wide increase in recombination rates compared to teosinte. The 12% increase in recombination rates results in ∼2 additional COs per meiosis, suggesting a tight upper bound on recombination rate evolution. Previous studies characterizing recombination rate variation between populations have concluded similarly: there is a limit to recombination rate evolution (67). However, it is possible that an increase greater than 12% would have become deleterious in maize and recombination rates can increase far beyond that, if the right conditions emerge. Nevertheless, if recombination rates increase too much, the abundance of chiasmata at the end of meiosis would cause improper chromosomal segregation, leading to sterility (69). As the main goal of most natural populations is to reproduce, tampering with meiosis could prove to have dire consequences.

### Maize-specific considerations for the evolution of the recombination landscape

It is known that post-domestication, maize hybridized with both teosintes – ssp. *parviglumis* and ssp. *mexicana* (70) resulting in gene flow between these populations. Migration and directional selection can favor increased recombination when selection at different loci covaries negatively between environments (18). These beneficial introgressions from *mexicana* were found in low recombining regions in maize that stayed low recombining after domestication (71) and therefore did not contribute beneficial cis-acting recombination-modifier alleles to maize but nevertheless could have created favorable conditions for recombination to evolve. Additionally, maize was domesticated in the lowlands of Mexico and had to adapt higher altitudes (72) which also might have created conditions that favored an increase in recombination.

Another consideration is that maize and teosinte possess chromosomal knobs, which are large regions of heterochromatin that suppress local recombination (60). Meiotic drive results from the interaction between these knobs and the meiotic drive locus on abnormal chromosome 10, Ab10, which causes preferential segregation of certain regions of the genome (73). Several studies have documented the differences between maize and teosinte knob occurrence and distribution (74); it was found that teosintes, *parviglumis* and *mexicana*, possess more interstitial knobs which could contribute to the observed suppression of recombination interstitially in teosinte. Nevertheless, knobs still occur in interstitial, gene-dense regions within maize so although knobs might be a factor in recombination landscape evolution during domestication, they are probably not solely responsible. Additionally, Ab10 is absent within modern inbred lines (75) so further testing if meiotic drive was responsible for the observed increase in recombination would have to be conducted in a landrace population.

Despite the above considerations, we believe their importance in the evolution of recombination landscape within maize to be small and instead we hypothesize that selection on domestication-related variation indirectly selected recombination modifiers involved in class I CO formation and CO interference. This process reduced CO interference on the longest chromosomes, allowing COs to be repositioned. The repositioning of COs was toward gene-dense regions that harbor domestication-related variation, shielding these regions from the Hill-Robertson effect. The benefits of altering the recombination landscape in maize outweighed the potential negatives, enabling selection to act more efficiently on domestication-related variation likely allowing maize to adapt much more quickly.

## Materials and Methods

### Experimental mapping populations & genetic map creation

To investigate the evolution of recombination rates in maize during domestication, we analyzed two experimental mapping populations. The first was the maize NAM population, consisting of 4,674 progeny and 93,000 crossover (CO) events. The second was an F2 population derived from crosses between wild-collected Palmar Chico *parviglumis* individuals, consisting of 3,052 progeny and 150,000 CO events (19–21). Both datasets underwent the same filtering process. We excluded individuals with more than 40 COs in the maize population and more than 30 COs in the teosinte population. For the maize NAM population, which was developed through an initial cross between B73, and a NAM parent followed by five generations of selfing, we anticipated a maximum of 40 detectable COs per individual due to the homozygosity generated from each sequential selfing reducing the ability to detect COs. For the teosinte F2 population, which involved only a single meiosis, we expected a maximum of 30 COs per individual. Additionally, we removed CO intervals exceeding 100kb because we cannot accurately determine the midpoint of the CO event because the location of the CO site within this interval created too much uncertainty. The teosinte population exhibited a high number of false-positive COs due to the Viterbi algorithm misidentifying structural variants as CO events because the teosinte population was aligned to the B73 reference genome, necessitating more filtering (21).

After applying these filters, we retained 41,000 COs for the maize population and 36,000 COs for the teosinte population. We then partitioned the genome into 2Mb bins and calculated the recombination rate for each bin and each population. Recombination rates were defined as the number of COs per interval divided by the number of individuals contributing COs, multiplied by 100, and normalized by the bin size in Mb. To obtain genetic map lengths, we calculated the cM distance between bins. We used: genetic position [current] = genetic position [previous] + (physical position[current] – physical position [previous]) *recombination rate[current] (34). To obtain genome-wide recombination rates, we simply divided the genetic map length of each chromosome by the physical length of the chromosome in Mb based on the B73v4 reference genome (29).

### Maize and teosinte populations used in ARG-inference

For ARG-inference, we used different individuals than those in the experimental populations mentioned above. ARG-inference relies upon DNA sequence variation and therefore we needed high quality sequenced individuals that were unrelated. We wanted individuals with whole genome sequencing (WGS) given WGS provides an unbiased and significant amount of information on the genome of interest. All individuals used in ARG-inference were sequenced using WGS.

The maize population consisted of 60 maize lines, including temperate, tropical, CIMMYT, and Chinese lines, randomly sampled from the 282 set, which was a panel of diverse maize lines constructed in the 1990s (30, 76). The VCF for the maize population was filtered using the maize HapMap3 tags; SNPs with ‘LLD’ tag, SNPs in LD with known GBS markers were retained while SNPs with ‘N15’ tag, SNPs < 5bp from known indels were removed (30). This filtering scheme yielded 31M SNPs genome-wide for the maize population. The teosinte population consisted of 50 *parviglumis* lines which were sampled from 5 distinct locations in Mexico: Amaltán de Cañas, Crucero Lagunitas, El Rodeo, Los Guajes, and San Lorenzo (31). The teosinte population was filtered identically to maize and yielded 30.5M SNPs genome wide. Before input into ARGweaver, the teosinte VCF was phased using Beagle software 5.4 (77). Both VCFs were aligned to B73v4 reference genome in previous studies.

### ARG-inference using ARGweaver software

To construct ARGs, we used the ARGweaver software (32) which relies on a Markov chain Monte Carlo (MCMC) algorithm to sample ARGs from a posterior distribution given the observed DNA sequence patterns and the model priors, such as the ratio between the recombination rate to mutation rate and N_e_ estimates going back in time. An example input into ARGweaver can be seen in our GitHub repository (https://github.com/ruthkepstein/Maize_recombination_evolution). ARGweaver is a Bayesian method and thus relies on several priors as input. First, we set effective population sizes (N_e_) at 0, 10,000, and 100,000 generations back in time to reflect the large population size of *Zea mays* pre-domestication, the domestication genetic bottleneck followed by exponential growth (14). We used similar N_e_ estimates in our teosinte population since it is known that teosinte went through a similar bottleneck during domestication (31). We also used a 2Mb recombination map as a prior, based on the maize NAM population and the F2 population of Palmar Chico teosintes mentioned above (25–27). Although our maize and teosinte populations used for ARG-inference were different than the populations used for the recombination map prior, the map had a resolution of 2Mb which is generalized and probably captures the broad-scaled patterns of the recombination landscapes. Because the recombination rate to mutation rate (ρ/µ) ratio must be 1 or greater to meet ARGweaver assumptions, we used the same recombination map as the mutation map to ensure each region had the ratio of 1. Genomic sites farther away than 5kb from genes were masked, as 95% of recombination events take place less than 5kb from genes, and because 85% of the maize genome is transposons which might create false signals within the ARG (25, 57). Lastly, to run ARGweaver efficiently, each chromosome was broken up into 1Mb segments and each segment was run on a separate CPU core to speed up inference. As well as parallelizing ARGweaver, we used a compression factor of 2 which takes every 2 invariant sites together to additionally shorten computational time. The posterior distribution of each 1Mb region was re-sampled for 1,000 Markov Chain Monte Carlo (MCMC) iterations; most 1Mb regions converged after 600 iterations, thus the first 600 iterations were discarded as burn-in. Since the teosinte individuals were collected from the wild and were heterozygous, we phased the teosinte individuals before input into ARGweaver and ran ARGweaver with —phased-vcf option. The maize VCF was derived from maize inbred lines, and thus these individuals should be homozygous at most sites. Instead of phasing, we used the first site of each diploid individual with the –-subsites option in ARGweaver.

Recombination rates are defined in ARGs as the number of recombination events per segment per MCMC iteration normalized by the branch length. After discarding the first 600 MCMC iterations as burn-in, we extracted recombination events and branch lengths at time zero. We utilized recombination events inferred at time zero, because Deng *et al.* 2021 observed a decline in inferred recombination rates as one traces further back into the ARG. This decline is attributed to ARGweaver’s limitation in detecting recombination events that do not alter the local tree structure, specifically when a lineage splits and immediately re-coalesces onto the same branch (78).

Because each MCMC iteration inferred recombination rates for differently sized segments, we normalized the recombination rates by dividing each inferred recombination rate by the length of the segment to obtain the recombination rate per 1bp. We then binned the genome into 100kb or 1Mb bins using the R package dplyr and calculated the mean recombination rate per bin. In addition to recombination rates, RTH values, were inferred from the ARG. We extracted RTH values using ARGweaver internal functions and averaged over MCMC iterations.

### Linear model construction to predict recombination rates using DNA methylation

We obtained ^m^CG and ^m^CHG data extracted from maize and teosinte individuals from a previous study (42). Using the ARG-inferred recombination rate intervals mentioned above, we annotated each 1Mb region with the mean DNA methylation level using bedmap software in command line. We then used the statsmodel package in Python to construct an ordinary least squares regression to predict recombination rates. We used a square root transformation of recombination rates to reduce heteroskedasticity, 10-fold cross-validation and centered the independent variables, by subtracting the mean from each observation, to avoid over-fitting of the model.

### Gene density correlation with recombination rates

Similar to the DNA methylation annotation of recombination rate intervals, we counted the number of genes per interval using the B73v4 reference annotation (29). We then correlated maize and teosinte recombination rates with gene density using the corr() function in Python. We used a Pearson correlation after transforming the recombination rates and gene density to achieve a normal distribution.

### Deleterious score dataset

We obtained zero-shot scores predicted for each SNP based on the 26 NAM parents (36). We calculated the mean zero-shot score per 1Mb interval, extracted intervals in the top 15% of recombination rate intervals for each population and compared their mean zero-shot score using a t-test.

### F_ST_ estimation

To estimate F_ST_, the maize and teosinte VCFs used in ARG-inference were combined using bcftools (79) and only overlapping SNPs were maintained. We conducted LD pruning on the combined VCF and removed SNPs with r^2^ > 0.3 in 10kb windows resulting in 776k SNPs genome wide. We then used vcftools (80) –-weir-fst-pop command to calculate F_ST_ at each SNP.

## Acknowledgements

We would like to thank Dr. Ed Buckler for his ARG insight and population genomic ideas, Dr. Jeff Ross-Ibarra for sharing his ideas and related thesis, Dr. April Wei for her help with understanding ARG creation, Dr. Jingjing Zhai and Dr. Wei-Yun Lai from the Buckler lab for creating the zero-shot score dataset, Erin Farmer and Quinn Simonis for their help with the linear models, and Dr. Gen Xu and lab for sharing their teosinte methylation data set with us.

