## Supplemental Tables & Figures for "The maize recombination landscape evolved during domestication"

### Supplementary Tables & Figures

**Supplementary Table 1.** Maize-specific model. $R^2 = 0.697$, F-statistic = 4242.

| | Coefficients | p-value |
|---|---|---|
| Maize $^{m}$CG % | -0.0894 | 0.000 *** |

**Supplementary Table 2.** Teosinte-specific model. $R^2 = 0.655$, F-statistic = 3502.

| | Coefficients | p-value |
|---|---|---|
| Teosinte $^{m}$CHG % | -0.0577 | 0.000 *** |

**Supplementary Table 3.** List of meiotic or recombination-related genes in the maize genome.

| Chr# | Start | End | Gene ID v5 | Gene name |
|---|---|---|---|---|
| chr1 | 2222930 | 2233674 | Zm00001eb000540 | *Prd4 / Prd3B* |
| chr1 | 9571476 | 9581461 | Zm00001eb003580 | *Flip* |
| chr1 | 26281990 | 26285480 | Zm00001eb008630 | *Rpa1* |
| chr1 | 43348941 | 43351082 | Zm00001eb013040 | *Top6A* |
| chr1 | 126548721 | 126590026 | Zm00001eb026880 | *Recq2* |
| chr1 | 157308972 | 157317169 | Zm00001d030741 | *Pam1* |
| chr1 | 180116252 | 180123158 | Zm00001eb032220 | *Swi1* |
| chr1 | 205851477 | 205868282 | Zm00001eb038490 | *Smc1* |
| chr1 | 228323641 | 228330878 | Zm00001eb043410 | *ERCC-1/ yeast Rad10* |
| chr1 | 234693720 | 234701386 | Zm00001eb045200 | *DNA LIGASE 1* |
| chr1 | 258031683 | 258046468 | Zm00001eb050670 | *MATERNAL EFFECT EMBRYO ARREST 70* |

| | | | | |
|---|---|---|---|---|
| chr1 | 258325654 | 258337156 | Zm00001eb050740 | *RecA1* |
| chr1 | 281342744 | 281347724 | Zm00001eb056730 | *XRCC4* |
| chr1 | 300801292 | 300803494 | Zm00001eb062810 | *JC73* |
| chr1 | 303550879 | 303557049 | Zm00001eb063880 | *Asy4a* |
| chr1 | 306411473 | 306420667 | Zm00001eb064930 | *Ku80* |
| chr10 | 1197798 | 1209464 | Zm00001eb405000 | *Smu2* |
| chr10 | 1746600 | 1750382 | Zm00001eb405340 | *Nse1* |
| chr10 | 15004735 | 15010645 | Zm00001eb408950 | *Mac1* |
| chr10 | 98434012 | 98465128 | Zm00001eb419130 | *Brca2* |
| chr10 | 124371838 | 124386615 | Zm00001eb423930 | *Zyp1* |
| chr10 | 127716085 | 127721835 | Zm00001eb424890 | *Pch2* |
| chr10 | 150798895 | 150807219 | Zm00001eb434000 | *Msh3* |
| chr2 | 5893895 | 5900903 | Zm00001eb068550 | *ESSENTIAL MEIOTIC ENDONUCLEASE 1B* |
| chr2 | 7132297 | 7137455 | Zm00001eb069270 | *Mre11a* |
| chr2 | 7871151 | 7879601 | Zm00001eb069600 | *Dv1 / divergent spindle 1* |
| chr2 | 11913362 | 11936797 | Zm00001eb071490 | *DNA ligase 4* |
| chr2 | 28059698 | 28070159 | Zm00001eb076670 | *CHROMATIN REMODELING 24* |
| chr2 | 30161987 | 30166229 | Zm00001eb077450 | *Brca1* |
| chr2 | 33010267 | 33019761 | Zm00001eb078170 | *PDS5 B-B* |
| chr2 | 65755582 | 65761280 | Zm00001eb084850 | *Mhf1* |
| chr2 | 160573693 | 160582923 | Zm00001eb094560 | *Mnd1* |
| chr2 | 198049825 | 198067735 | Zm00001eb102420 | *As1/ Asy1* |
| chr2 | 200535133 | 200542866 | Zm00001eb103090 | *Gen1* |
| chr2 | 204187046 | 204205675 | Zm00001eb104340 | *Mlh3* |

| | | | | |
|---|---|---|---|---|
| chr2 | 204273249 | 204295539 | Zm00001eb104360 | *Mlh3* |
| chr2 | 206251501 | 206303314 | Zm00001eb105070 | *Msh4* |
| chr2 | 232784991 | 232798172 | Zm00001eb114240 | *Rfc1* |
| chr2 | 239339326 | 239352162 | Zm00001eb116810 | *Ku70* |
| chr2 | 240922104 | 240939102 | Zm00001eb117500 | *Facnd2* |
| chr3 | 3164814 | 3170508 | Zm00001eb119810 | *Acoz1* |
| chr3 | 9360039 | 9375366 | Zm00001eb122180 | *Mus2* |
| chr3 | 35921449 | 35923982 | Zm00001eb127650 | *Rpa3B* |
| chr3 | 38829415 | 38838043 | Zm00001eb128110 | *Sun2* |
| chr3 | 97372242 | 97407571 | Zm00001eb133210 | *Figl1* |
| chr3 | 137067752 | 137071229 | Zm00001eb138190 | *Rad51A2* |
| chr3 | 153165414 | 153169795 | Zm00001eb141350 | *Mus81-1* |
| chr3 | 153817250 | 153821621 | Zm00001eb141480 | *RecA3* |
| chr3 | 170672154 | 170682606 | Zm00001eb144310 | *Smc2* |
| chr3 | 173727756 | 173732411 | Zm00001eb145140 | *Zip4* |
| chr3 | 179418627 | 179429367 | Zm00001eb146450 | *XRCC2* |
| chr3 | 196592651 | 196601719 | Zm00001eb151620 | *Exo1A* |
| chr3 | 225740978 | 225745413 | Zm00001eb160270 | *Rad51c* |
| chr3 | 235455861 | 235460581 | Zm00001eb163340 | *Dmc1* |
| chr4 | 3366098 | 3370929 | Zm00001eb165270 | *RecqI1* |
| chr4 | 31840297 | 31846067 | Zm00001eb171920 | *Mre11b* |
| chr4 | 34756199 | 34759975 | Zm00001eb172550 | *Spo11-2* |
| chr4 | 93551464 | 93568073 | Zm00001eb180360 | *Prd2* |
| chr4 | 104252297 | 104337711 | Zm00001eb181120 | *Rad50* |
| chr4 | 133996051 | 134020836 | Zm00001eb183500 | *Pms1* |

| | | | | |
|---|---|---|---|---|
| chr4 | 145143457 | 145171139 | Zm00001eb184840 | *Mer3* |
| chr4 | 188955616 | 188981213 | Zm00001eb195020 | *Dfo* |
| chr4 | 212955137 | 212983699 | Zm00001eb201140 | *Fancm1* |
| chr4 | 247070262 | 247075093 | Zm00001eb208520 | *Cdc1* |
| chr5 | 7553723 | 7558287 | Zm00001eb214790 | *Spo11-1* |
| chr5 | 11990381 | 11995625 | Zm00001eb216450 | *Hop2* |
| chr5 | 16714877 | 16726059 | Zm00001eb218470 | *Am1* |
| chr5 | 18091348 | 18097149 | Zm00001eb218750 | *MATERNAL EFFECT EMBRYO ARREST 7* |
| chr5 | 24281844 | 24315496 | Zm00001eb220610 | *DNA mismatch protein MutS type 2* |
| chr5 | 27340966 | 27345552 | Zm00001eb221260 | *Nbs1* |
| chr5 | 60884322 | 60890572 | Zm00001eb227420 | *Mtopvib1* |
| chr5 | 75134860 | 75152580 | Zm00001eb230860 | *Invan6* |
| chr5 | 79351074 | 79364575 | Zm00001eb231860 | *Recq4* |
| chr5 | 90720078 | 90729450 | Zm00001eb233650 | *Sun1* |
| chr5 | 91735021 | 91741995 | Zm00001eb233800 | *Asy3 / Dsy2* |
| chr5 | 153121542 | 153131785 | Zm00001eb239370 | *Hei10* |
| chr5 | 166057998 | 166061646 | Zm00001eb241470 | *DNA helicase* |
| chr5 | 177373803 | 177375013 | Zm00001eb243950 | *Xrcc3* |
| chr5 | 183840644 | 183849294 | Zm00001eb245990 | *BRCA1-associated RING domain* |
| chr5 | 191711783 | 191719193 | Zm00001eb248070 | *RPA2* |
| chr5 | 193085552 | 193107390 | Zm00001eb248620 | *Shoc1* |
| chr5 | 210017470 | 210033528 | Zm00001eb252400 | *Recg1* |
| chr5 | 218116420 | 218134744 | Zm00001eb255780 | *CHR25, HOMOLOG OF RAD54* |

| | | | | |
|---|---|---|---|---|
| chr5 | 225047534 | 225050107 | Zm00001eb259380 | *RPA2* |
| chr6 | 10128299 | 10132839 | Zm00001eb260930 | *Kin1* |
| chr6 | 14745661 | 14753594 | Zm00001eb261800 | *Tdm1* |
| chr6 | 82512206 | 82541538 | Zm00001eb270700 | *RadA-like / RecA family profile* |
| chr6 | 122235227 | 122277208 | Zm00001eb279300 | *Mcm9* |
| chr6 | 130465241 | 130497449 | Zm00001eb280900 | *Psd5A* |
| chr6 | 137588485 | 137612510 | Zm00001eb282720 | *Tubg3* |
| chr6 | 167825722 | 167838974 | Zm00001eb291710 | *Smc4* |
| chr6 | 169580675 | 169586236 | Zm00001eb292490 | *Brca1* |
| chr6 | 178086613 | 178093423 | Zm00001eb297030 | *Afd1 / Rec8* |
| chr6 | 179510358 | 179529022 | Zm00001eb297550 | *Smc3* |
| chr7 | 2019511 | 2041340 | Zm00001eb298600 | *Scc2* |
| chr7 | 19036353 | 19043975 | Zm00001eb303090 | *Sgo1* |
| chr7 | 22199838 | 22201533 | Zm00001eb303730 | *HIGLE / SLX1 homolog* |
| chr7 | 27750867 | 27756831 | Zm00001eb304600 | *Asy4b* |
| chr7 | 147241326 | 147249342 | Zm00001eb318880 | *Gen2* |
| chr7 | 169205732 | 169212956 | Zm00001eb325310 | *Rad51A1* |
| chr7 | 171840364 | 171850168 | Zm00001eb326520 | *Mus1* |
| chr7 | 174150168 | 174156859 | Zm00001eb327280 | *Spc97 / Spc98* |
| chr7 | 179106476 | 179111901 | Zm00001eb329200 | *Rad51d* |
| chr8 | 78200600 | 78206647 | Zm00001eb345090 | *Ptd* |
| chr8 | 124103394 | 124114076 | Zm00001eb352660 | *Msh5* |
| chr8 | 124138401 | 124156551 | Zm00001eb352670 | *Msh5* |
| chr8 | 163577828 | 163587939 | Zm00001eb362590 | *Mlh1* |
| chr8 | 173783809 | 173811580 | Zm00001eb367170 | *Asp1* |

| | | | | |
|---|---|---|---|---|
| chr9 | 54860680 | 54881073 | Zm00001eb382430 | *Phs1* |
| chr9 | 106616254 | 106619181 | Zm00001eb388310 | *Com1* |
| chr9 | 115330275 | 115341438 | Zm00001eb389920 | *Prd1* |
| chr9 | 151718656 | 151725634 | Zm00001eb399650 | *SDS* |
| chr9 | 153220844 | 153224509 | Zm00001eb400180 | *Rfa1* |
| chr9 | 161337579 | 161345236 | Zm00001eb404350 | *Prd3* |

**Supplementary Table 4.** List of maize lines used to construct maize ARG.

| | | |
|---|---|---|
| 282set_CML258 | 282set_A679 | 282set_Mo24W |
| 282set_NC344 | 282set_ILLHy | 282set_CML281 |
| 282set_NC368 | 282set_KY228 | 282set_Ci7Goodman-Buckler |
| 282set_Pa91 | 282set_Tzi10 | 282set_B105 |
| 282set_Ms71 | 282set_NC250 | 282set_NC340 |
| 282set_N28Ht | 282set_B73 | 282set_Yu796 |
| 282set_A635 | 282set_SA24 | 282set_B104 |
| 282set_R229 | 282set_Ab28A | 282set_SC213R |
| 282set_K64 | 282set_B68 | 282set_CM37 |
| 282set_CML321 | 282set_NC328 | 282set_NC294 |
| 282set_NC318 | 282set_A441-5 | 282set_NC236 |
| 282set_KY226 | 282set_CML314 | 282set_CML333 |
| 282set_NC352 | 282set_Os420 | 282set_W182B |
| 282set_Va59 | 282set_NC346 | 282set_B52 |
| 282set_NC264 | 282set_CH9 | 282set_C49A |
| 282set_CI66 | 282set_Oh40B | 282set_E2558W |
| 282set_CML103 | 282set_Va26 | 282set_Mp339 |
| 282set_CML322 | 282set_38-11Goodman-Buckler | 282set_D940Y |
| 282set_GT112 | 282set_SC55 | 282set_CML258 |
| 282set_NC320 | 282set_CML254 | |
| 282set_R168 | 282set_NC358 | |

**Supplementary Table 5.** List of teosinte individuals used in teosinte ARG.

| | |
|---|---|
| FS1821_02 | FS1886_08 |
| FS1821_03 | FS1886_10 |
| FS1821_04 | FS1886_11 |
| FS1821_05 | FS1886_12 |
| FS1821_06 | FS1904_01 |
| FS1821_07 | FS1904_02 |
| FS1821_08 | FS1904_03 |
| FS1821_09 | FS1904_04 |
| FS1821_10 | FS1904_06 |
| FS1838_13 | FS1904_07 |
| FS1838_14 | FS1904_08 |
| FS1838_15 | FS1904_09 |
| FS1838_16 | FS1904_10 |
| FS1838_17 | FS1904_11 |
| FS1838_18 | FS1837_01 |
| FS1838_19 | FS1837_02 |
| FS1838_20 | FS1837_05 |
| FS1838_21 | FS1837_08 |
| FS1838_22 | FS1837_09 |
| FS1886_01 | FS1837_10 |
| FS1886_02 | FS1837_11 |
| FS1886_03 | FS1837_12 |
| FS1886_04 | FS1837_03 |
| FS1886_05 | FS1821_02 |
| FS1886_06 | FS1821_03 |

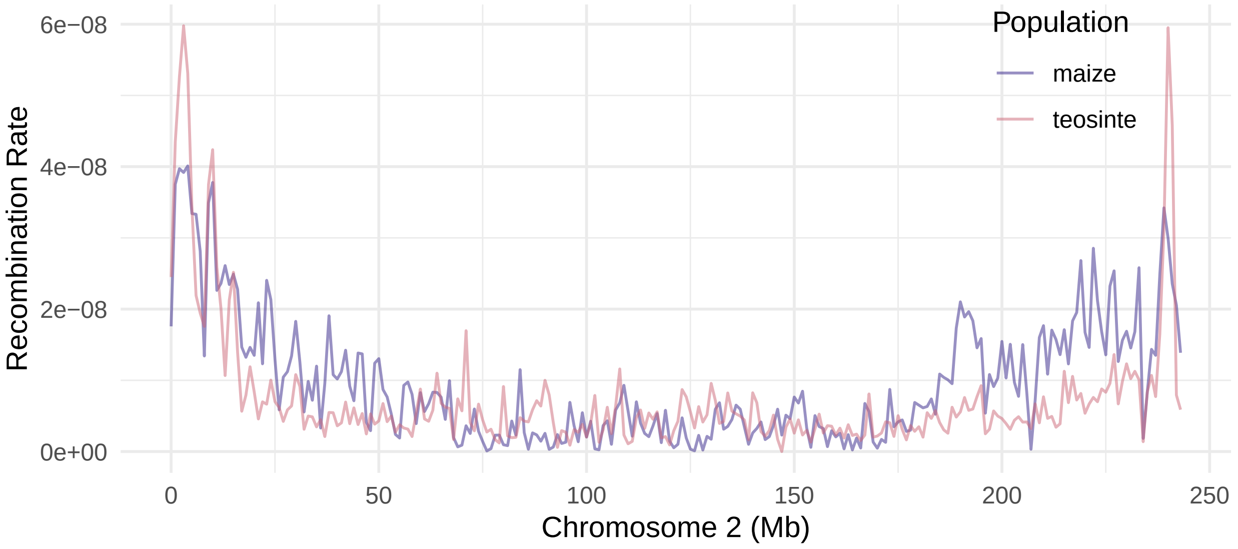
Population
maize
teosinte
Recombination Rate
6e−08
4e−08
2e−08
0e+00
0
50
100
150
200
250
Chromosome 2 (Mb)

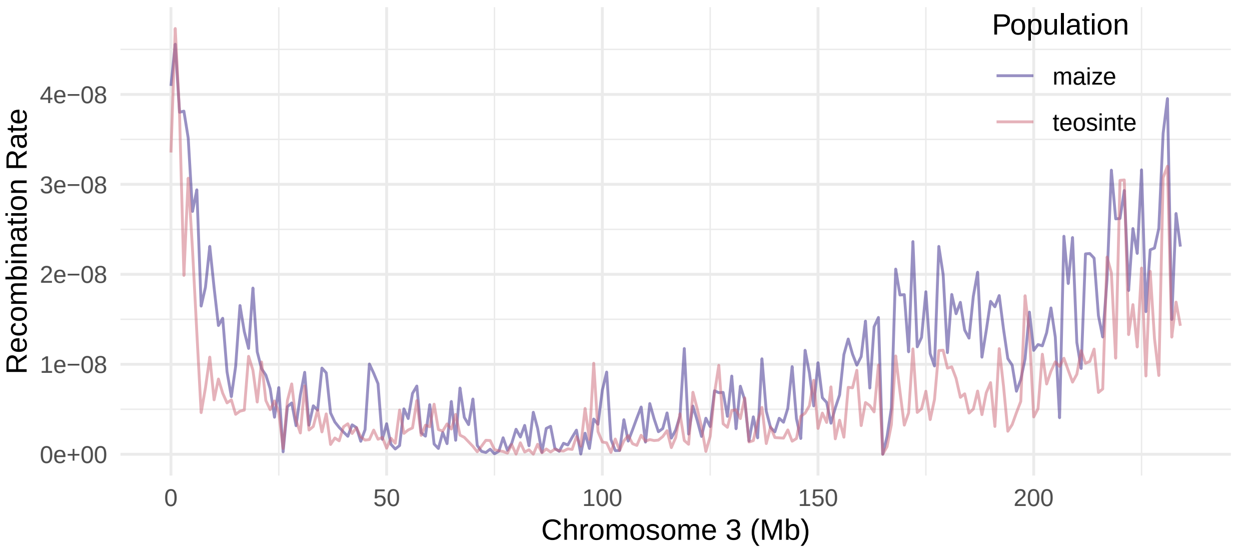
Population
maize
teosinte
Recombination Rate
4e−08
3e−08
2e−08
1e−08
0e+00
0
50
100
150
200
Chromosome 3 (Mb)

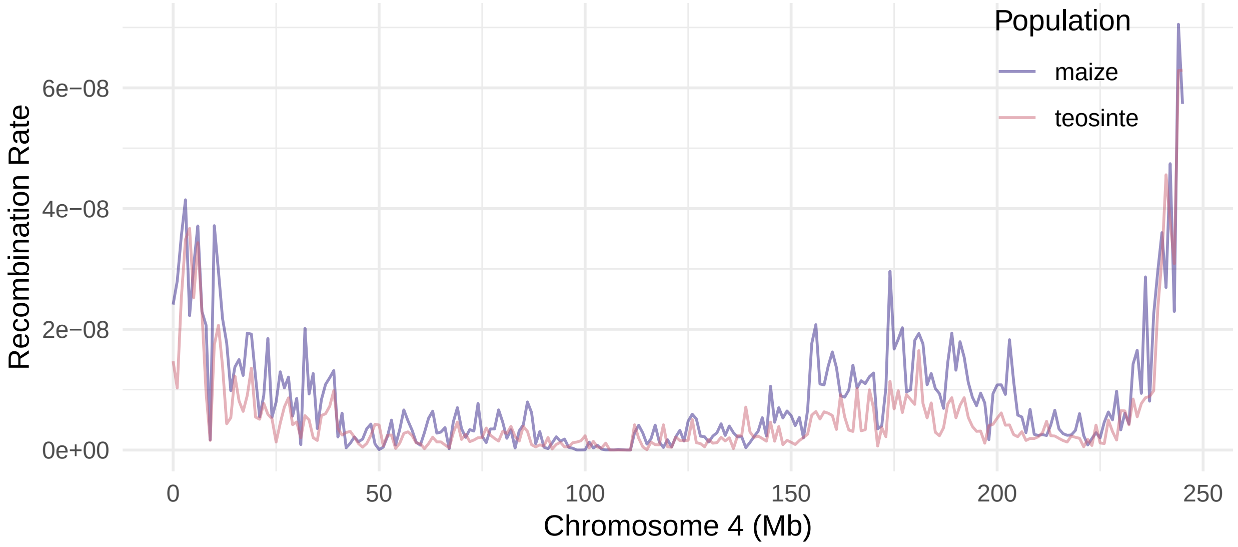
Population
maize
teosinte
Recombination Rate
6e−08
4e−08
2e−08
0e+00
0
50
100
150
200
250
Chromosome 4 (Mb)

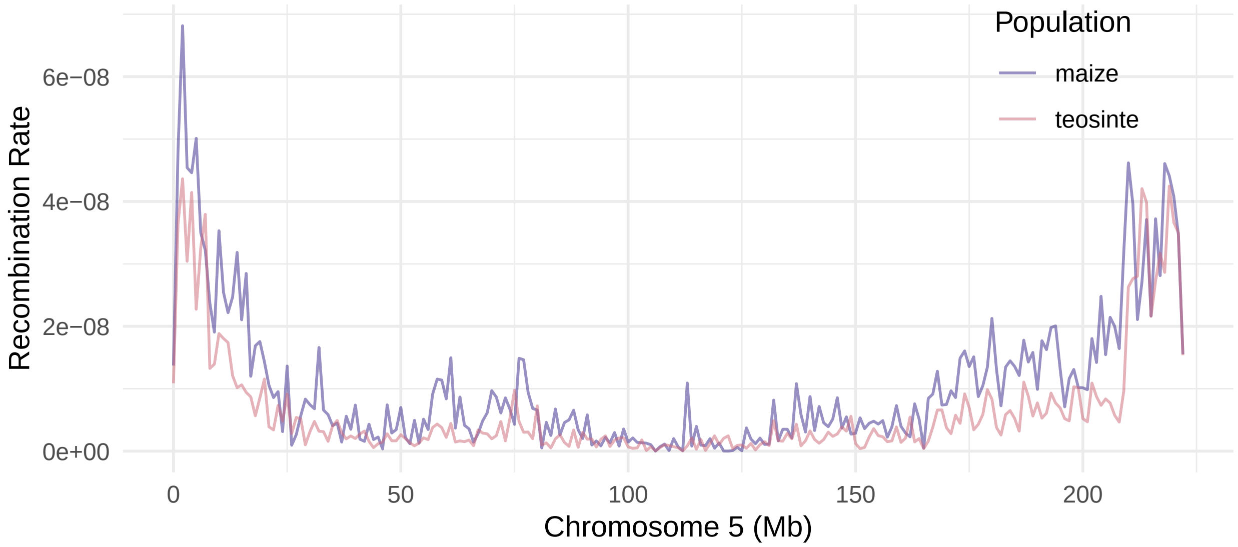
Population
maize
teosinte
Recombination Rate
6e−08
4e−08
2e−08
0e+00
0
50
100
150
200
Chromosome 5 (Mb)

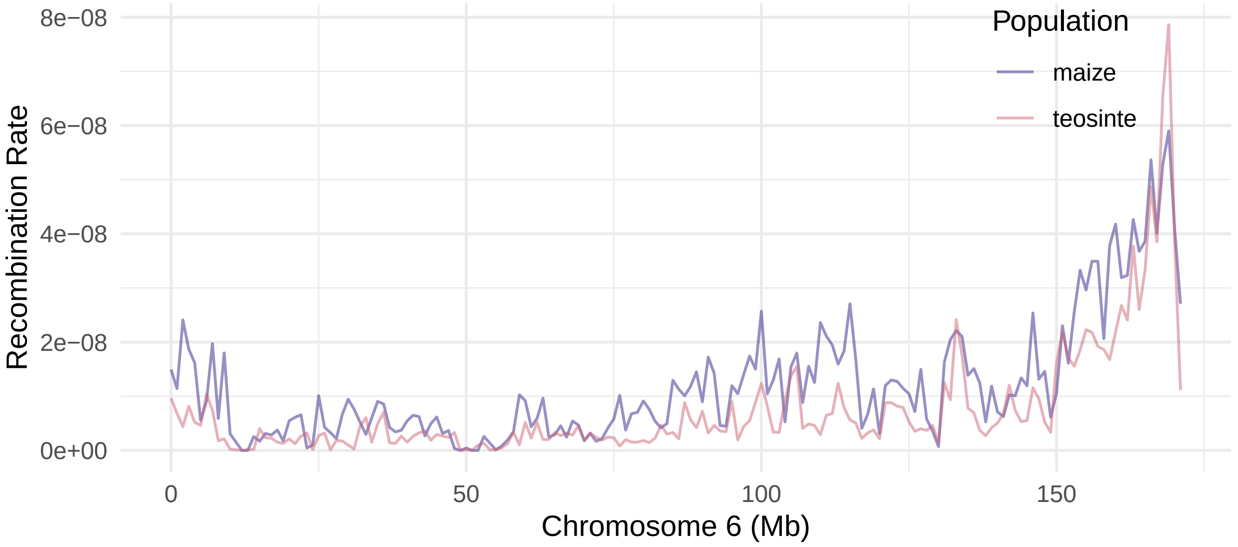
Population
maize
teosinte
Recombination Rate
8e−08
6e−08
4e−08
2e−08
0e+00
0
50
100
150
Chromosome 6 (Mb)

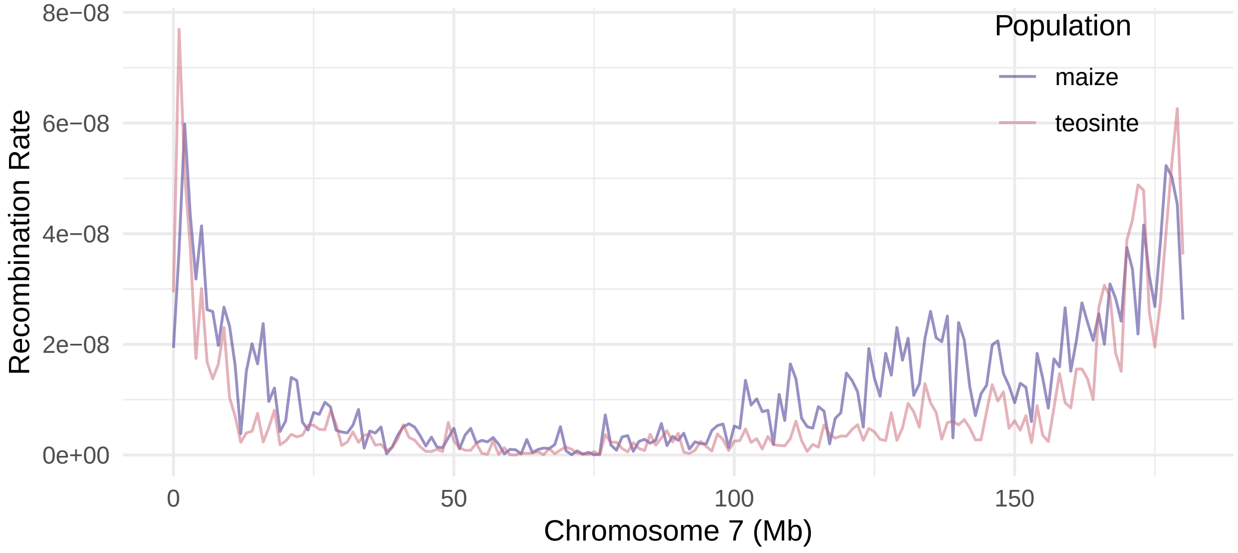
Population
maize
teosinte
Recombination Rate
8e−08
6e−08
4e−08
2e−08
0e+00
0
50
100
150
Chromosome 7 (Mb)

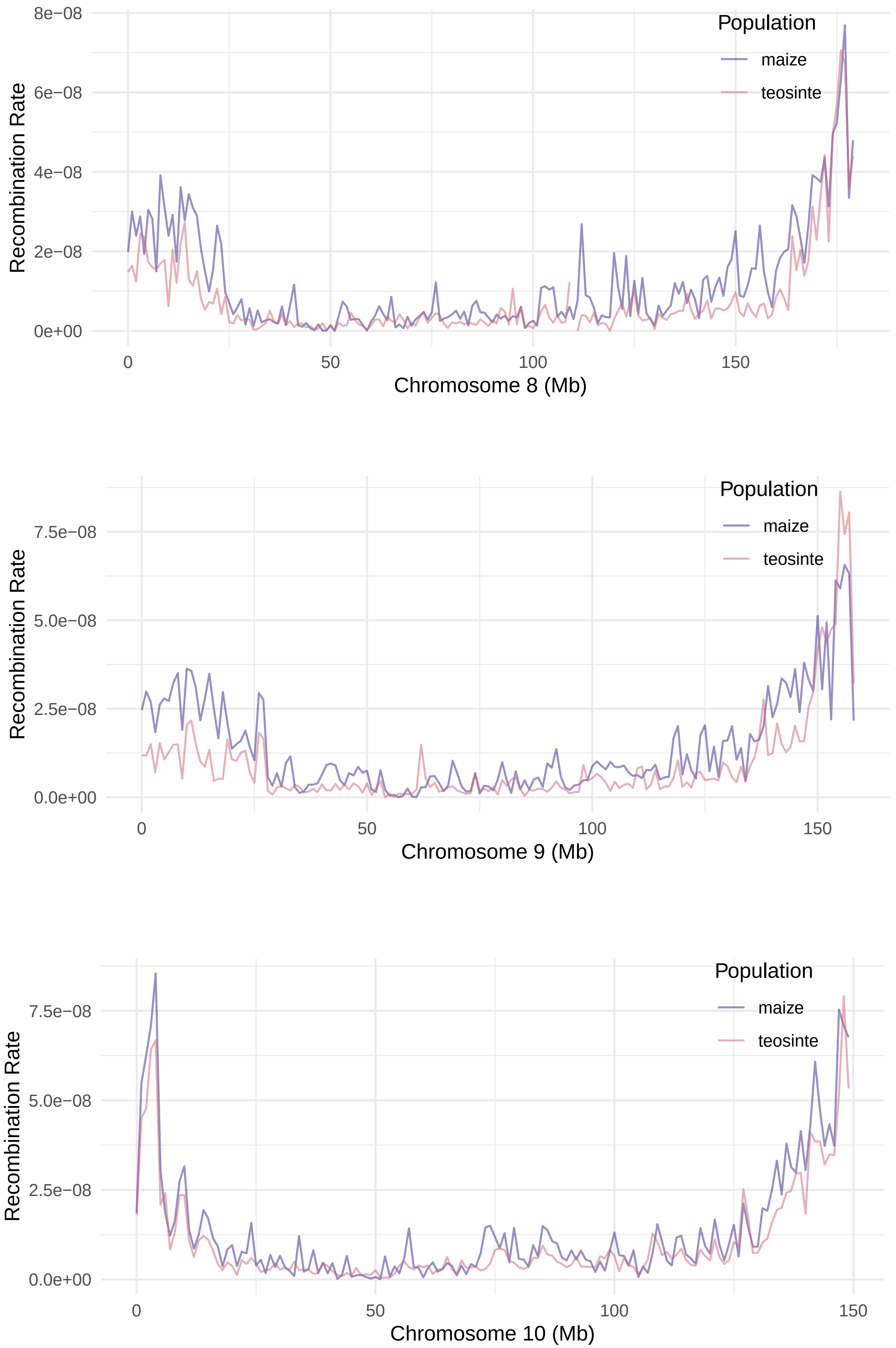


**Supplementary Figure 1.** ARG-inferred recombination rates for maize and teosinte across chromosomes 2-10.

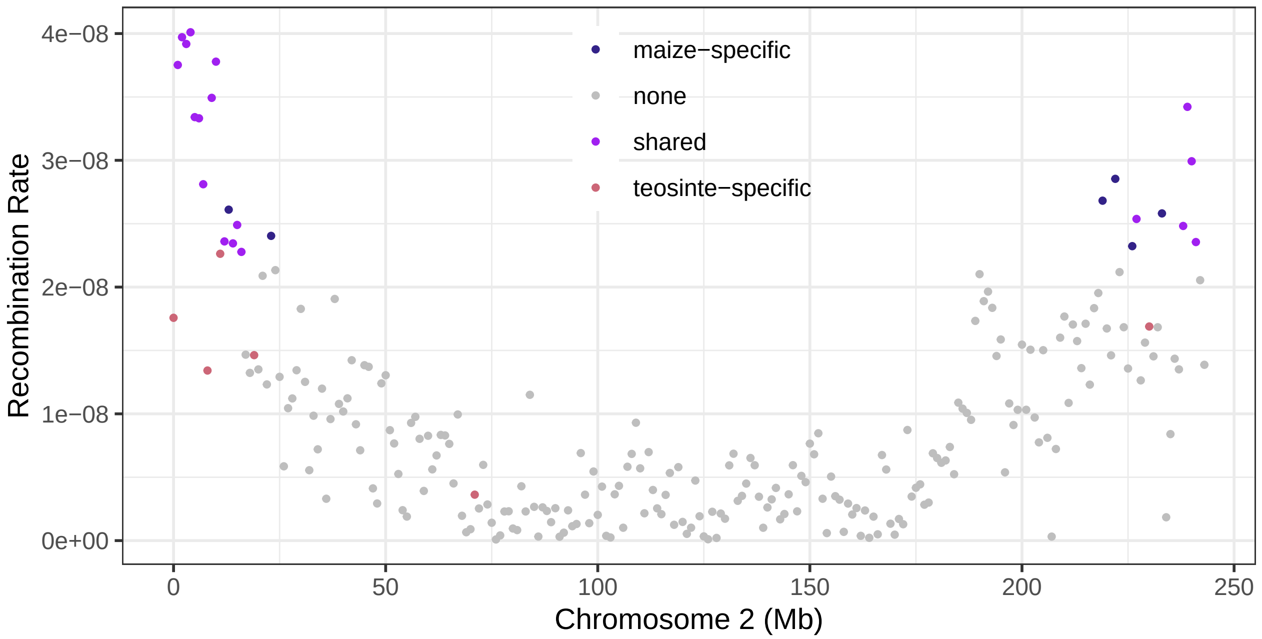
maize−specific
none
shared
teosinte−specific
Recombination Rate
4e−08
3e−08
2e−08
1e−08
0e+00
0
50
100
150
200
250
Chromosome 2 (Mb)

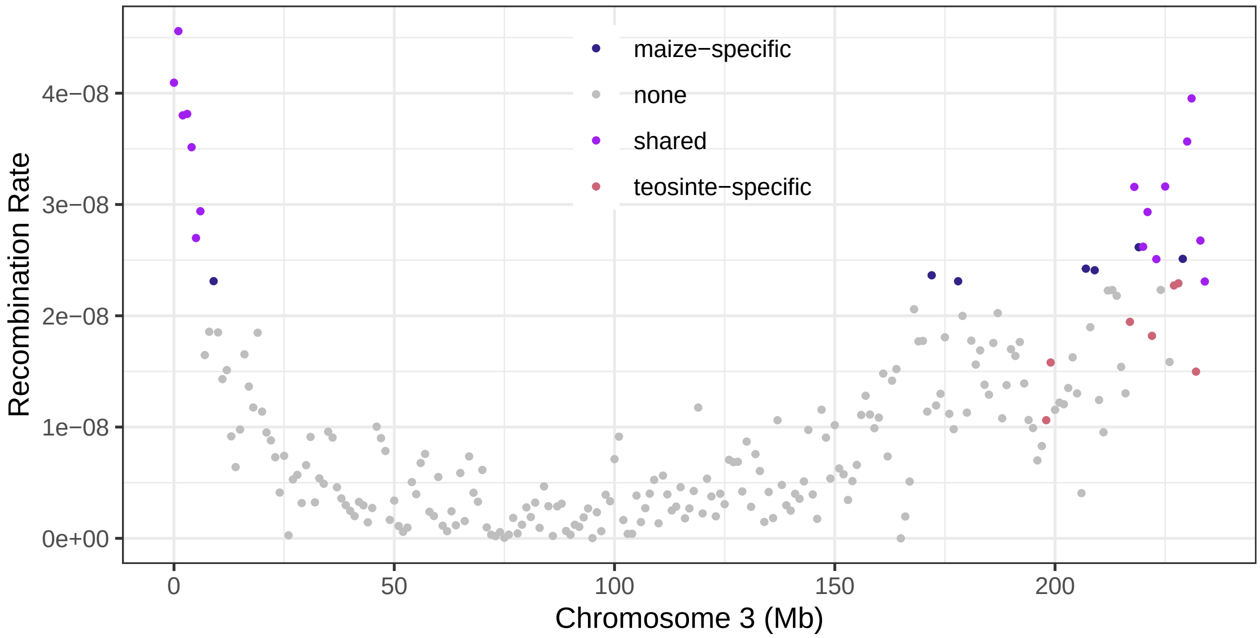
maize−specific
none
shared
teosinte−specific
Recombination Rate
4e−08
3e−08
2e−08
1e−08
0e+00
0
50
100
150
200
Chromosome 3 (Mb)

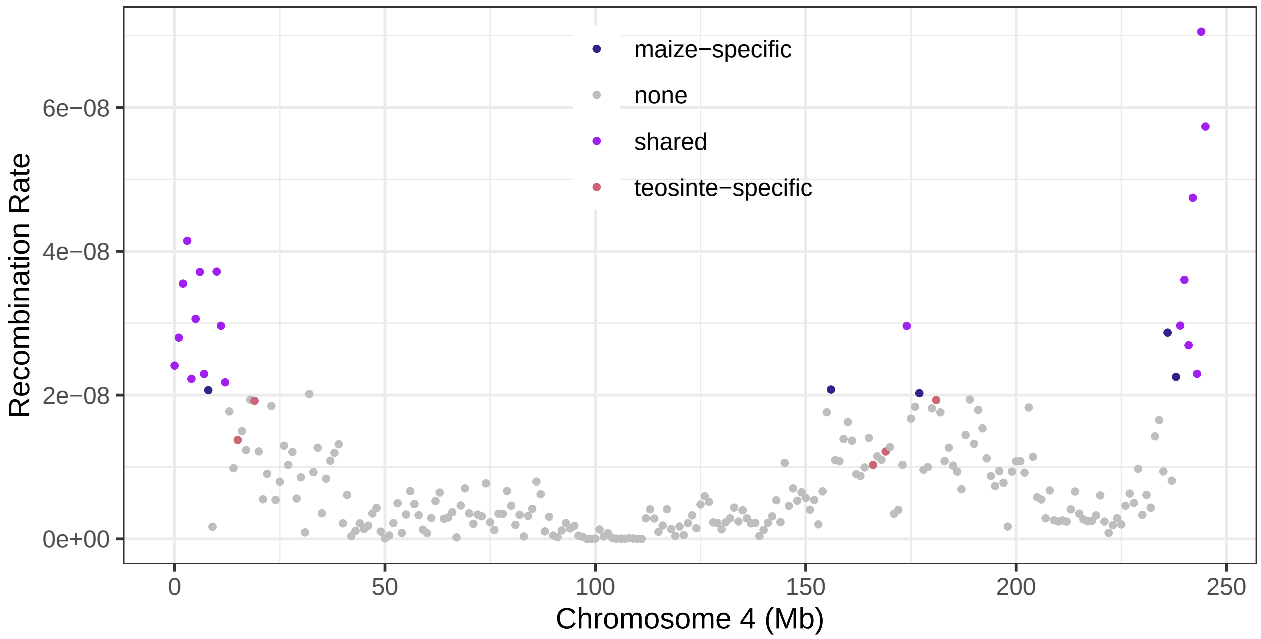
maize−specific
none
shared
teosinte−specific
Recombination Rate
6e−08
4e−08
2e−08
0e+00
0
50
100
150
200
250
Chromosome 4 (Mb)

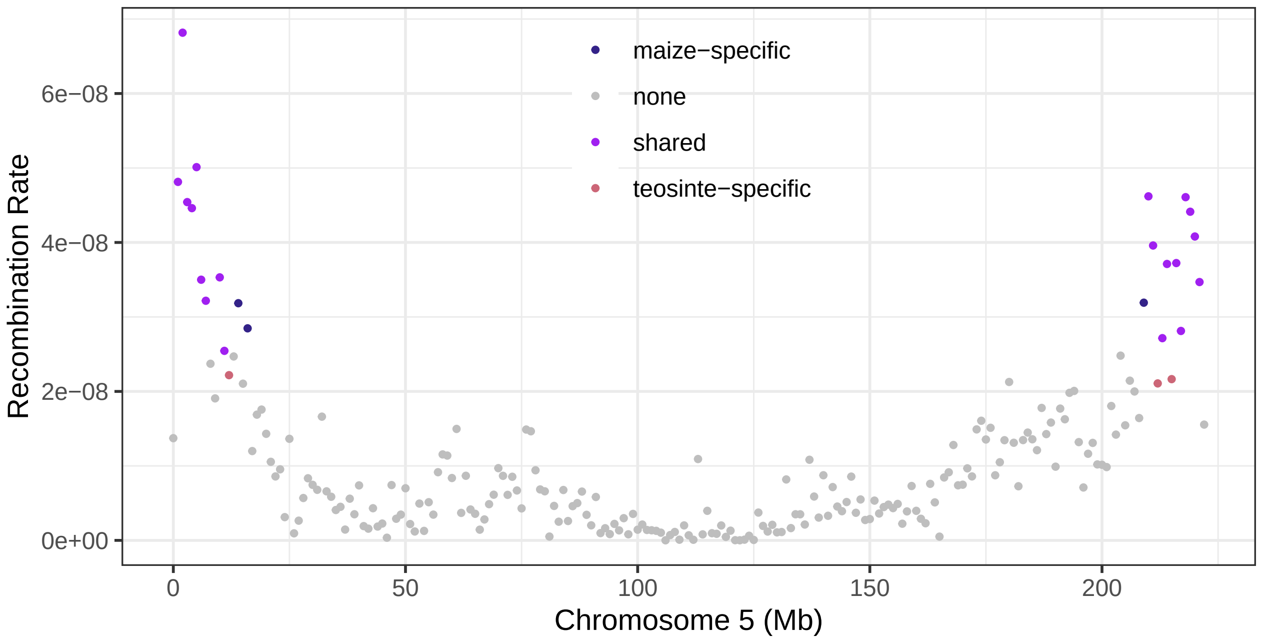
maize−specific
none
shared
teosinte−specific
Recombination Rate
6e−08
4e−08
2e−08
0e+00
0
50
100
150
200
Chromosome 5 (Mb)

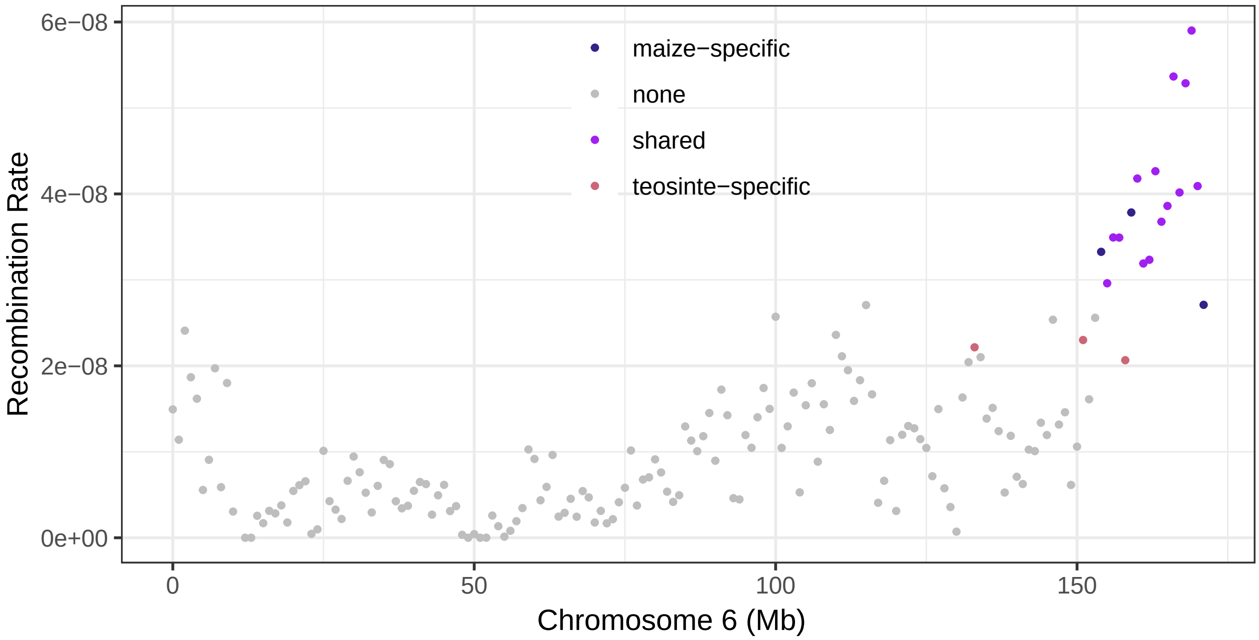
maize−specific
none
shared
teosinte−specific
Recombination Rate
6e−08
4e−08
2e−08
0e+00
0
50
100
150
Chromosome 6 (Mb)

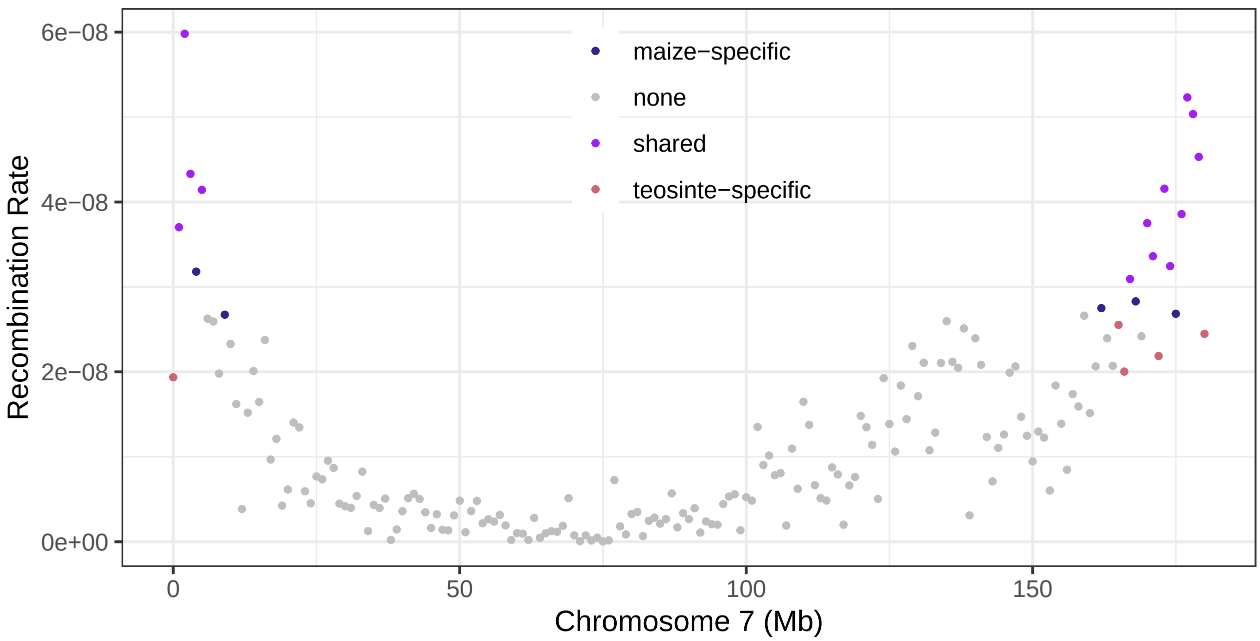
maize−specific
none
shared
teosinte−specific
Recombination Rate
6e−08
4e−08
2e−08
0e+00
0
50
100
150
Chromosome 7 (Mb)

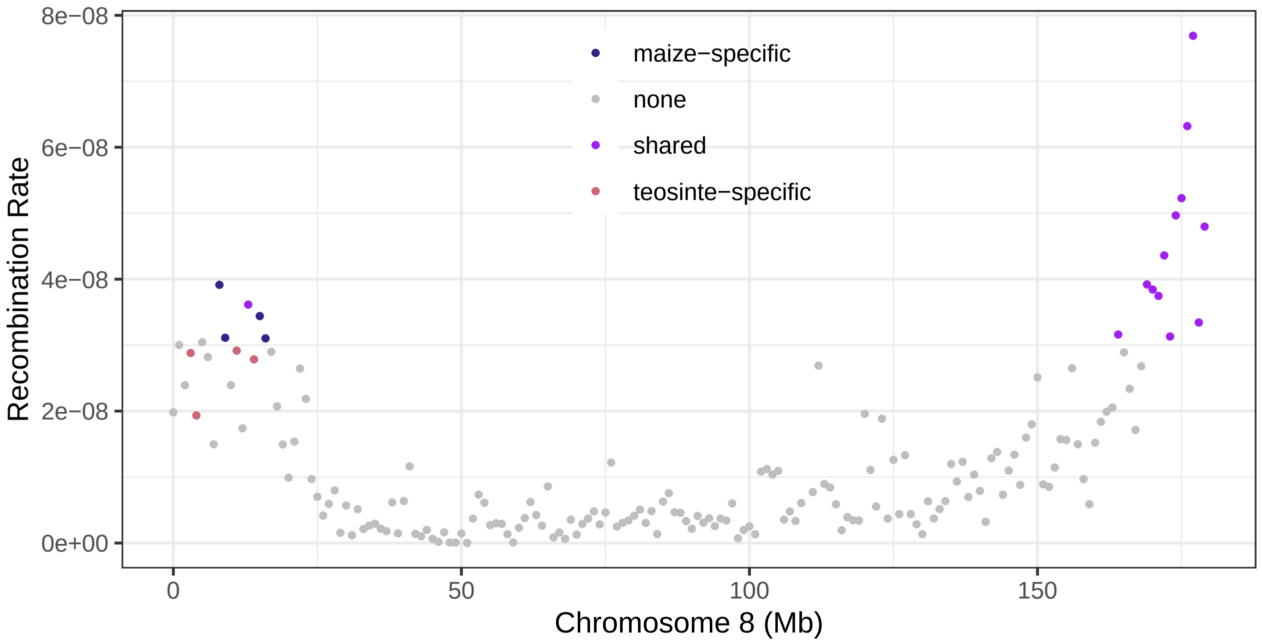


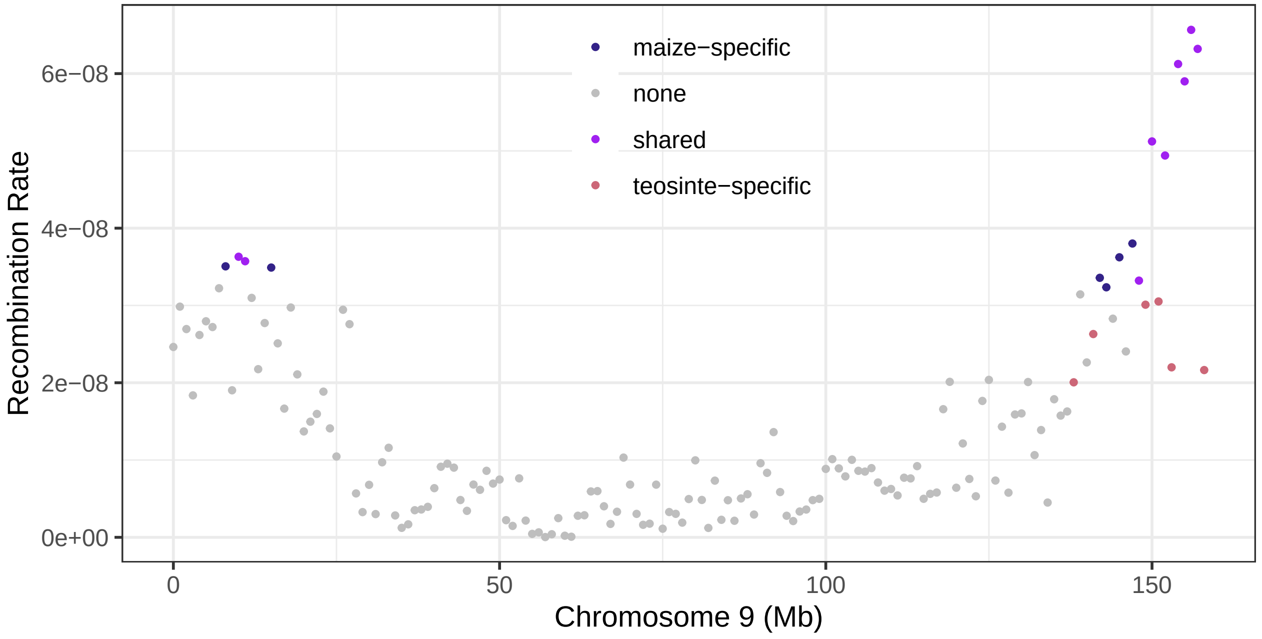


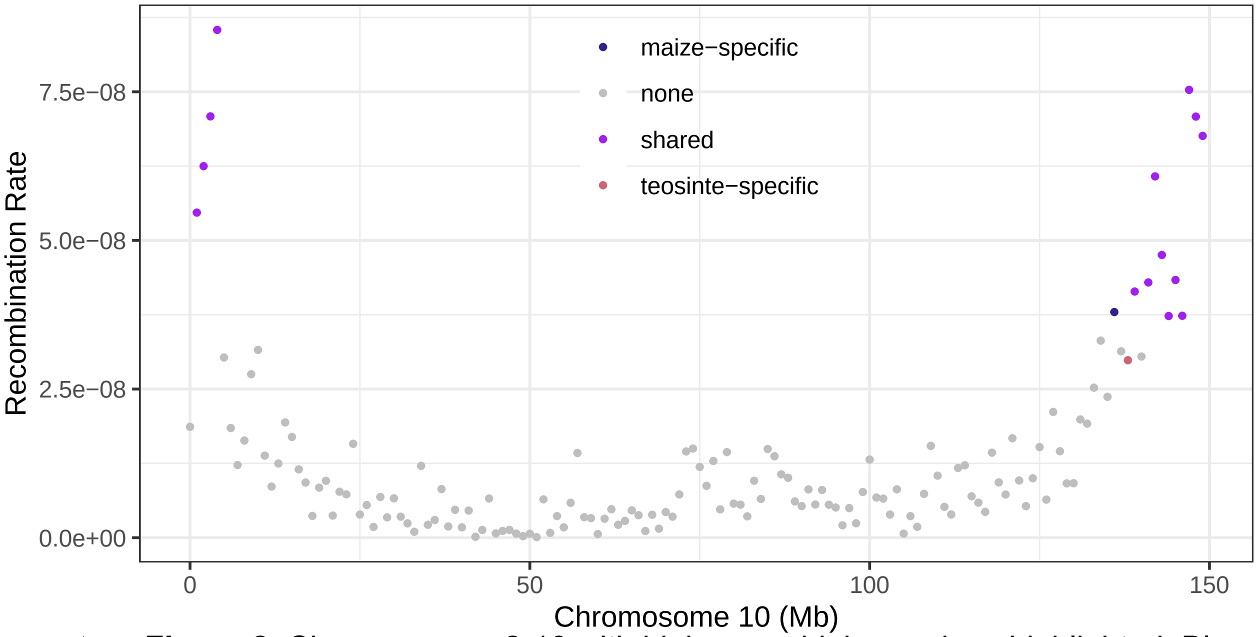


**Supplementary Figure 2.** Chromosomes 2-10 with high recombining regions highlighted. Blue = maize high recombining regions, Pink = teosinte high recombining regions, and Purple = shared high recombining regions.

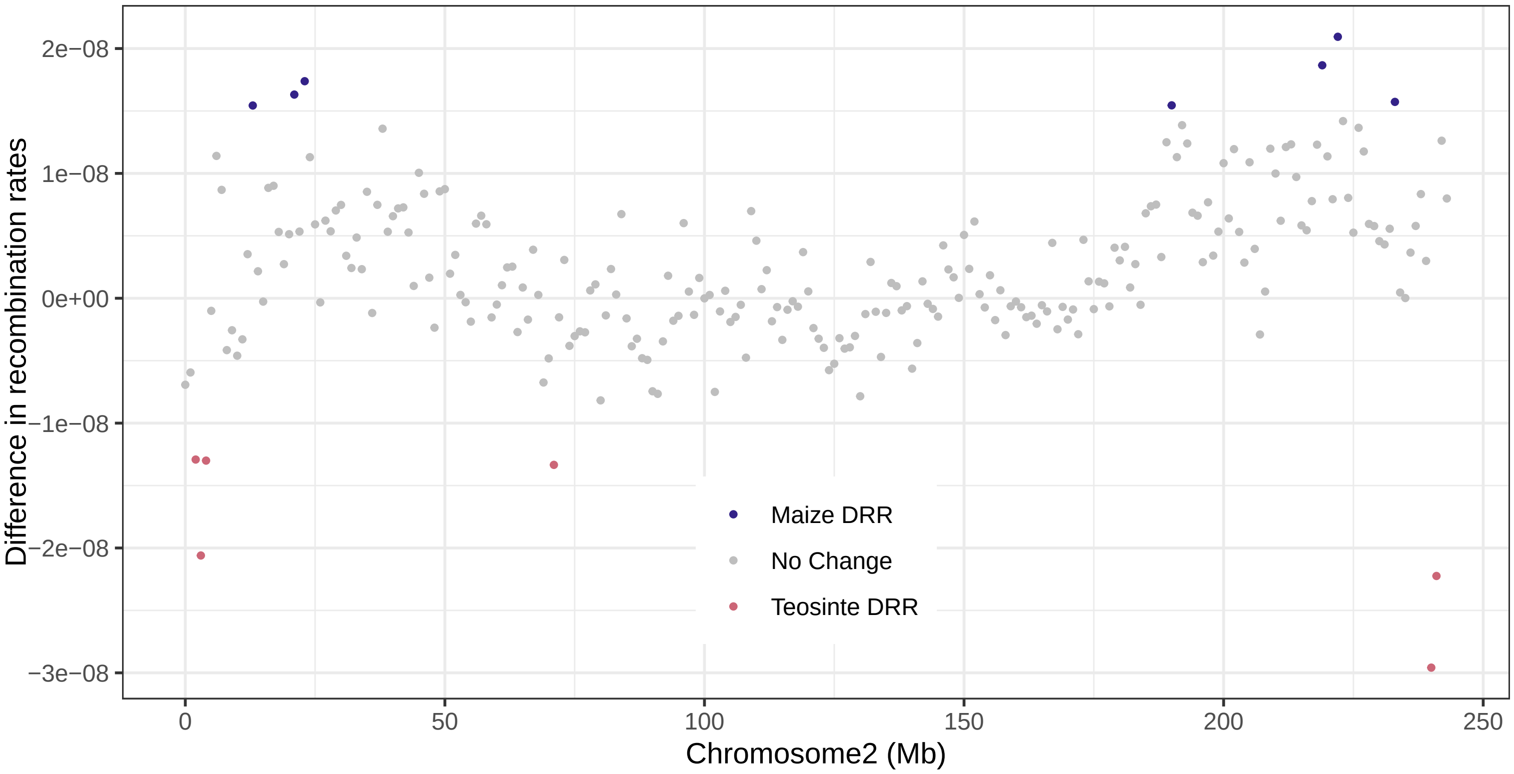
Difference in recombination rates
2e−08
1e−08
0e+00
−1e−08
−2e−08
−3e−08
Maize DRR
No Change
Teosinte DRR
0
50
100
150
200
250
Chromosome2 (Mb)

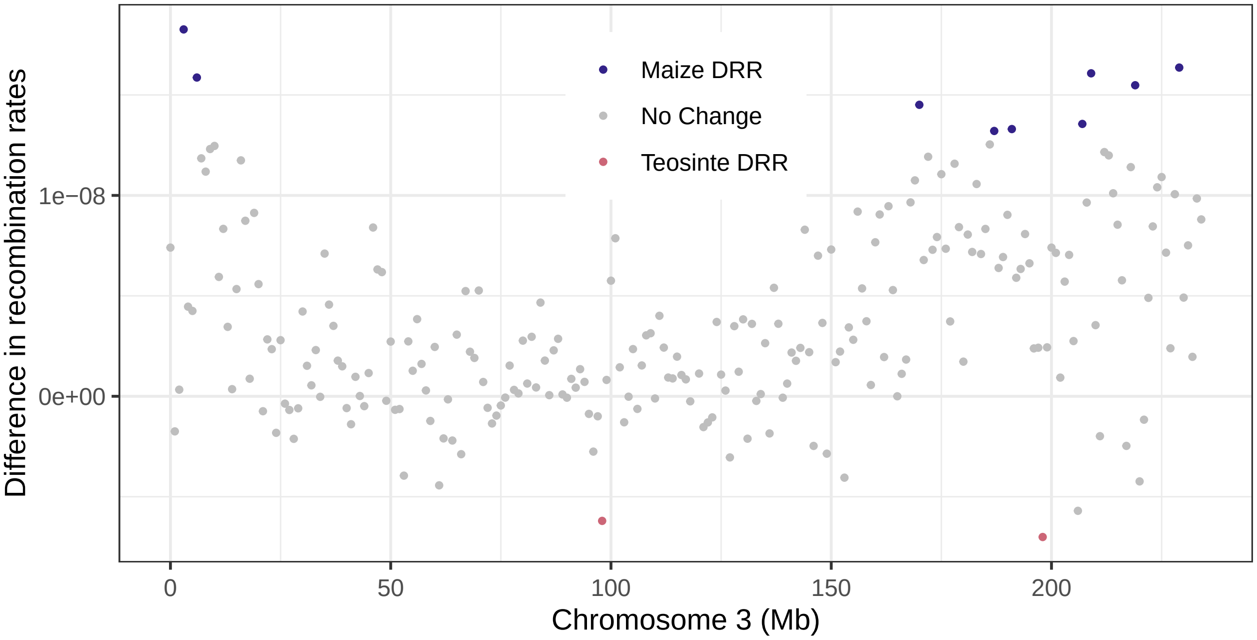
Difference in recombination rates
1e−08
0e+00
Maize DRR
No Change
Teosinte DRR
0
50
100
150
200
Chromosome 3 (Mb)

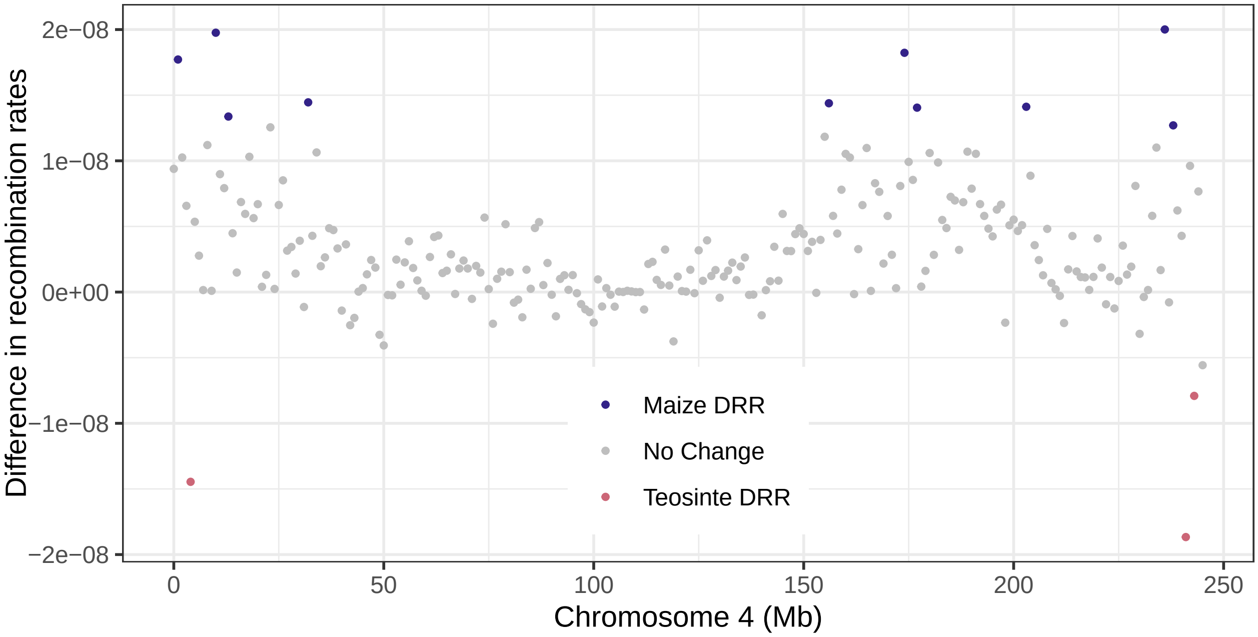
Difference in recombination rates
2e−08
1e−08
0e+00
−1e−08
−2e−08
Maize DRR
No Change
Teosinte DRR
0
50
100
150
200
250
Chromosome 4 (Mb)

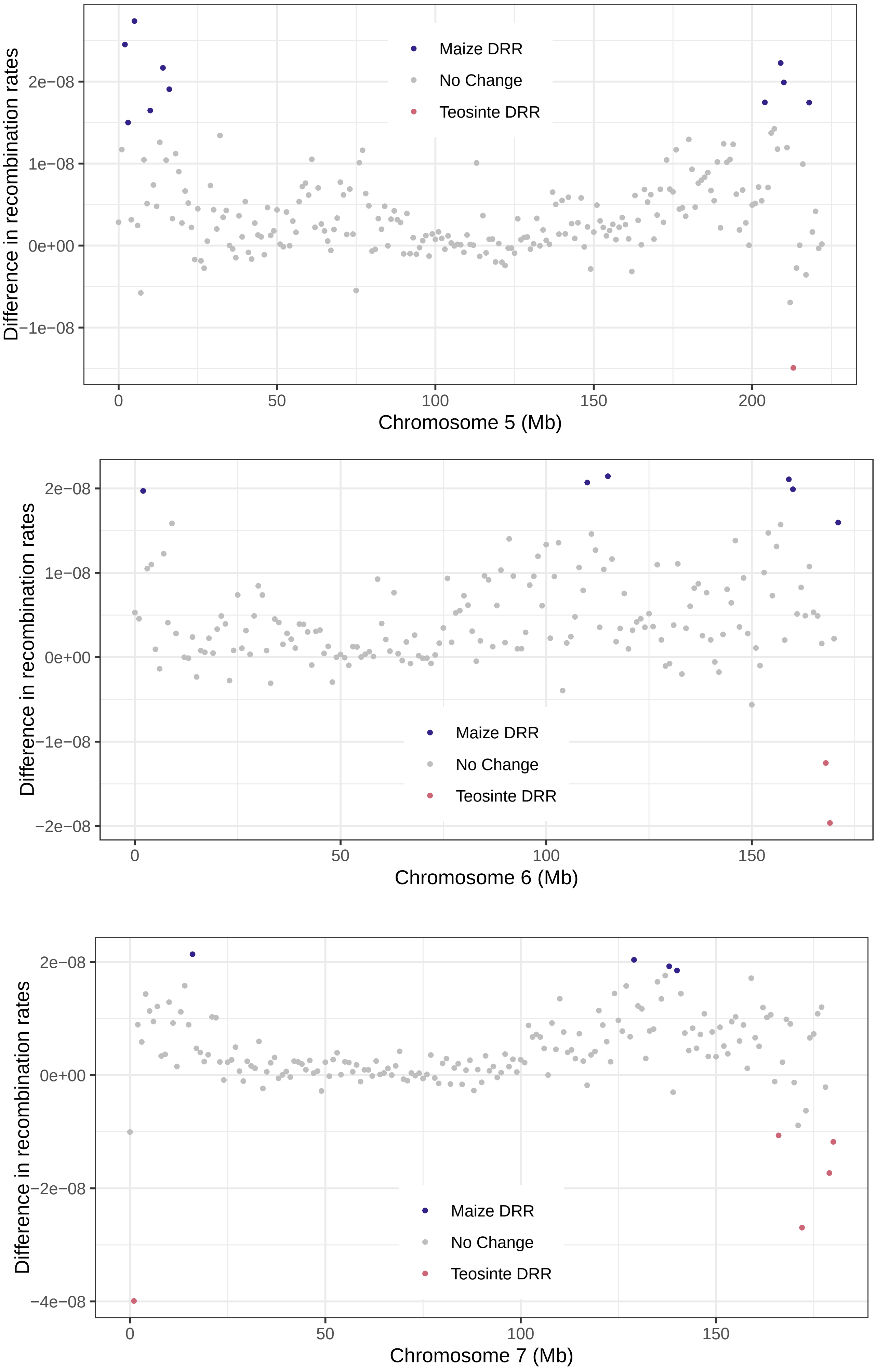

Maize DRR
No Change
Teosinte DRR
Difference in recombination rates
2e−08
1e−08
0e+00
−1e−08
0
50
100
150
200
Chromosome 5 (Mb)
Maize DRR
No Change
Teosinte DRR
Difference in recombination rates
2e−08
1e−08
0e+00
−1e−08
−2e−08
0
50
100
150
Chromosome 6 (Mb)
Maize DRR
No Change
Teosinte DRR
Difference in recombination rates
2e−08
0e+00
−2e−08
−4e−08
0
50
100
150
Chromosome 7 (Mb)

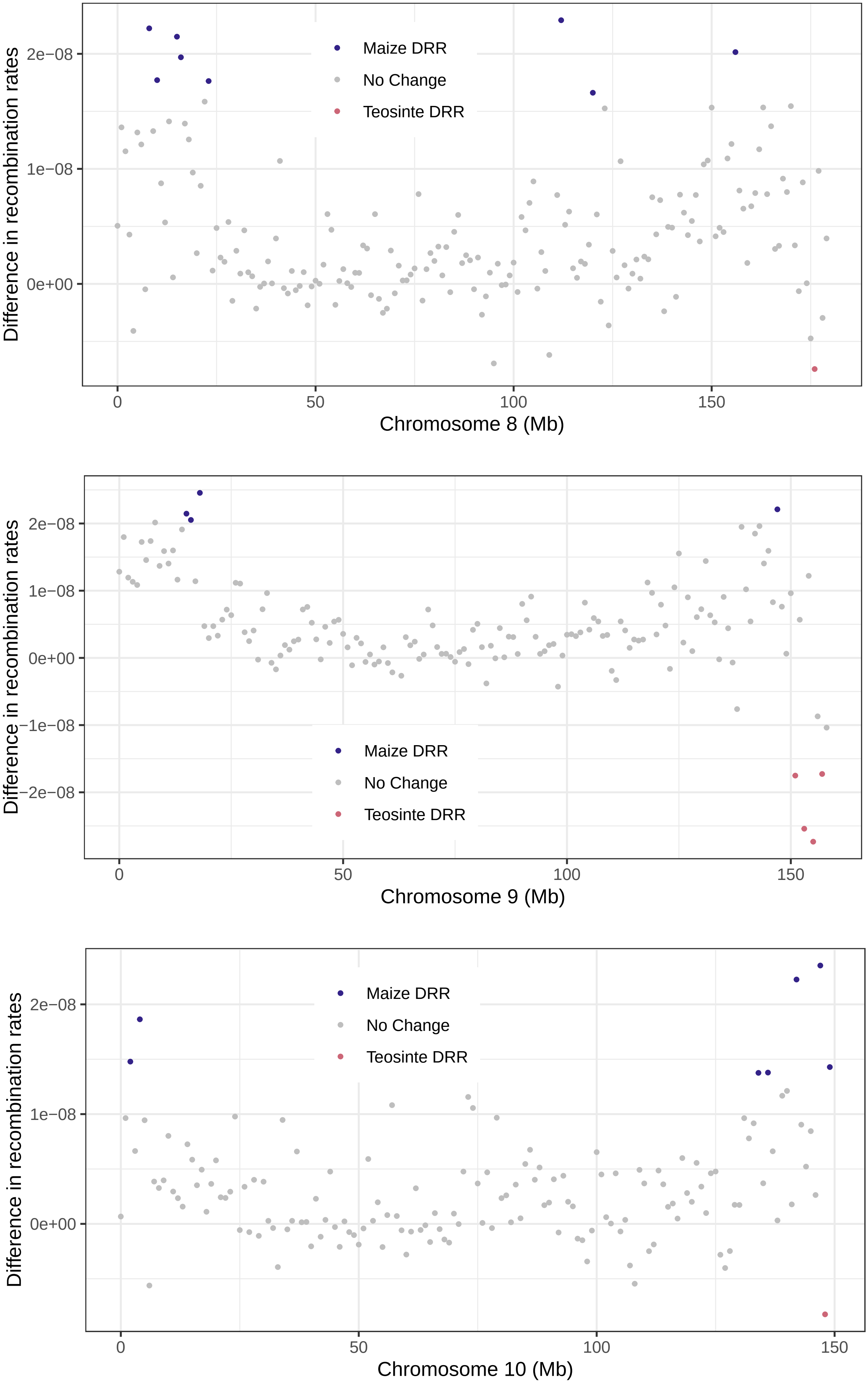


**Supplementary Figure 3**. Maize and Teosinte DRRs on chromosomes 2-10. Blue = maize DRR, Pink = teosinte DRR, grey = no significant difference in recombination.

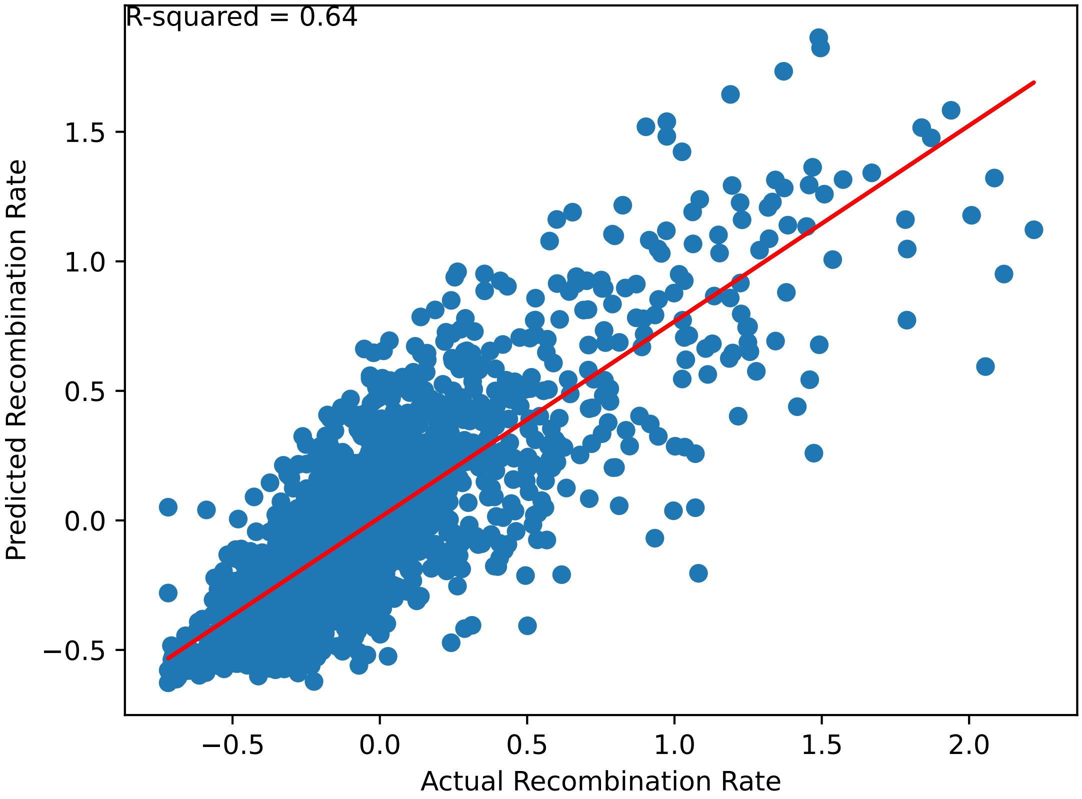


**Supplementary Figure 4.** Predicted teosinte recombination rates vs. actual teosinte recombination rates using the maize model trained on maize-specific data.

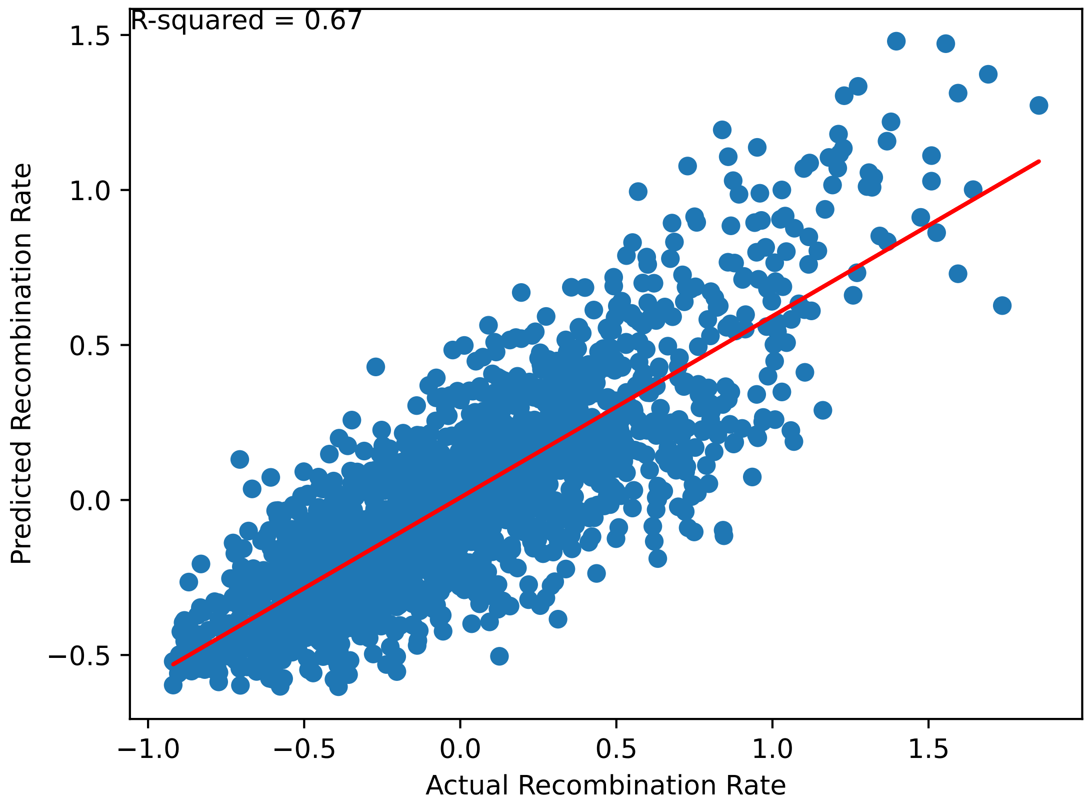


**Supplementary Figure 5.** Predicted maize recombination rates vs. actual maize recombination rates using the teosinte model trained on teosinte-specific data.
